# Organosulfur cycling in the multi-partner symbiosis between lucinid bivalves, sulphur-oxidizing symbionts, and *Endozoicomonas*

**DOI:** 10.64898/2025.12.05.692423

**Authors:** Marta Sudo, Chuang Sun, Ornella Carrión, Omaima Zaki, Olivier Gros, Jay Osvatic, Joana Séneca, Jonathan D. Todd, Jillian M. Petersen

## Abstract

Organosulfur compounds play an important role in chemotaxis, stress protection, and nutrition in many marine symbioses. Lucinidae, an ancient and species-rich family of marine bivalves, host chemosynthetic sulphur-oxidising bacteria within gill epithelial cells, and is well-known for its contribution to inorganic sulphur cycling in marine sediments worldwide. Little is known about organosulfur transformations by lucinid-associated microbiota or the host’s potential contribution. Here, we integrated field measurements, genomics, transcriptomics, and experimental assays to investigate organosulfur cycling in three lucinid species from temperate and tropical environments. Concentrations of the osmolyte dimethylsulphoniopropionate (DMSP) in lucinids were typically much higher than in their surrounding environment, and varied with host species and season, but not with symbiont abundance. Indeed, candidate DMSP synthesis *DSYB* genes were expressed by the animal hosts implying that the host may be the source of DMSP. Furthermore, a newly identified and isolated *Candidatus* Endozoicomonas endolucinida symbiont was capable of DMSP catabolism to dimethylsulphide (DMS), which could be further syntrophically oxidised by the sulphur-oxidising symbionts that encode methanethiol oxidase genes in their genomes. Finally, incubation experiments supported a combined role of the host and its associated microbiome in organosulfur cycling during stress, with the highest DMSP concentrations measured when holobionts were incubated with antibiotics under anoxic conditions. This study identified a previously unrecognised microbial partner in lucinid symbioses and revealed its potential organosulphur-based metabolic interactions with both the host and the sulphur-oxidising symbionts, highlighting the capacity of these widespread associations to influence global organosulfur cycling.

## Introduction

Dimethylsulfoniopropionate (DMSP) is one of Earths most abundant organosulfur molecules, with global production exceeding 10⁹ tons annually ^1^. It is ubiquitous in marine ecosystems and mediates key biological and biogeochemical processes, from salinity and oxidative stress protection to microbial signalling, sulphur cycling, and climate regulation ^2–7^. Importantly, DMSP is prevalent in marine photosymbiotic systems, such as corals ^8^, anemones ^9^, and giant clams ^10^. Indeed, coral reefs are well-known hotspots of DMSP production and cycling, and the coral host itself was the first animal shown to produce DMSP *de novo* ^11–13^, with most animals containing DMSP previously thought to acquire it from their diet or phototrophic symbionts ^9^. Within the coral holobiont, both the photosynthetic symbionts (*Symbiodiniaceae*) ^14,15^ and associated bacteria can produce and cycle DMSP ^16,17^.

DMSP catabolism is a major source of carbon and sulphur for marine microorganisms ^18^, particularly in marine animal-microbe interactions ^8,19–21^. Such catabolism can yield climate-active gases including methanethiol (MeSH) and dimethylsulphide (DMS), thereby bridging marine and atmospheric sulphur cycles and significantly impacting atmospheric chemistry and global climate ^22–24^. DMSP can be demethylated via *dmd* gene products in bacteria for roles in C and S assimilation. This pathway is thought to predominate in the environment and can yield MeSH ^25^. Alternatively, diverse microbes and corals themselves can cleave DMSP to liberate dimethyl sulphide (DMS) plus a three-carbon compound, usually acrylate, via *ddd* or *Alma* gene products ^20,25–27^. DMSP cleavage can be utilised for C assimilation, respiration and/or chemotaxis ^5,8,19–21,28^. Note, there are diverse Alma-family DMSP lyase enzymes in algae and corals, plus nine distinct Ddd enzymes in diverse bacteria and fungi ^29^, including many associated with marine animal hosts, e.g., as *Pseudomonas* ^20^ and *Endozoicomonas* ^21^. *Endozoicomonas* is prevalent in corals worldwide and is associated with various other marine animals^30^, including sea slugs ^31^, sponges ^32^, sea anemones ^33^, scallops ^34^, and lucinid clams that host chemosynthetic sulphur-oxidizing symbionts ^35^. Reflecting this broad host range and the substantial genomic and metabolic diversity observed among species, *Endozoicomonas* has been proposed to occupy a mutualism-parasitism continuum, with putative functions that vary across hosts and environments^30,36,37^. In corals, *Endozoicomonas* is thought to support host health^38,39^, for example, by degrading DMSP and thereby reducing signals that attract pathogens such as *Vibrio coralliilyticus*, which uses DMSP to locate stressed hosts ^28,40,41^. A similar potentially beneficial interaction was recently described in the scallop *Argopecten irradians*. This scallop accumulates high levels of DMSP ^42^ and hosts endosymbiotic bacteria, which break down DMSP using the DMSP lyase (DddP) ^43^.

Further symbiotic animals with obvious potential for active organosulfur cycling are clams from the lucinidae family, which rely on their sulphur-oxidizing gammaproteobacterial endosymbionts for nutrition ^44^. The energy produced by sulphur oxidation is coupled with inorganic carbon fixation, which provides nutrition for the symbionts and the host ^44–47^. Intriguingly, genes predicted to encode subunits of dimethyl sulfoxide (DMSO) reductase, that produces DMS, are some of the highest expressed by lucinid symbionts, but there is so far no experimental demonstration of organosulfur cycling by these symbionts ^48^. Furthermore, while lucinid holobionts are distributed globally across diverse environments, their greatest diversity is found in shallow-water habitats such as seagrass meadows known to contain far higher DMSP levels than the overlying seawater ^49,50^. Although lucinid clams are usually described as two-partner associations comprised of the host and often only one species of sulphuroxidizing symbionts, recent studies have revealed unexpected complexity in the lucinidassociated microbiome, reporting several co-occurring sulphur-oxidizing symbiont strains or species, plus other associated bacteria, including *Endozoicomonas* ^35,51^.

Despite its importance in marine ecosystems and holobiont health in other symbioses, neither DMSP, nor its catabolites, have been studied in marine sulphur-oxidising symbioses. Given the global prevalence of lucinid holobionts in coastal sediments, their contribution represents a key missing link to understanding the coupling of inorganic and organosulfur cycles and their potential influence on environment and climate. Here, we combine field observations, multiomics, and experimental assays to investigate the potential contribution of lucinid hosts and associated microbes (*Ca*. Thiodiazotropha and *Endozoicomonas*) to DMSP synthesis and its degradation. Our study establishes lucinid holobionts as dynamic hotspots of organosulfur cycling in seagrass sediments and highlights their overlooked role in coastal sulphur fluxes.

## Methods

### Sample collection

Lucinid clams were sampled during four occasions: August 2021, November 2021, January 2022, and August 2022, from Piran, Slovenia, and Guadeloupe (Lesser Antilles). In Piran, samples were collected at “Bernardin Plaza” (45.5158279 N, 13.5764505 E) and “Italian Border” (45.5918590 N, 13.7161088 E). *Loripes orbiculatus* and *Loripinus fragilis* were collected from *Cymodocea nodosa* seagrass meadows via sediment extraction and sieving, with immediate preparation for DMSP measurements (see below). Species were identified based on morphological characteristics. In November 2021, 15 individuals of *L. orbiculatus* were dissected into (a) gills and (b) bodies containing visceral mass, mantle, and foot, with tissues from different individuals pooled to obtain sufficient biomass, resulting in two samples for DMSP measurements. In January 2022, 23 *L. orbiculatus* and 52 *L. fragilis* individuals were sampled, resulting in three gill and four body samples. In August 2022, ten *L. orbiculatus* and seven *L. fragilis* were collected, resulting in three samples for each species. Due to their small size, body and gill samples of *L. fragilis* were combined, which resulted in inconclusive data and were excluded from further analysis. During this sampling, triplicate DMSP samples of sediment and *C. nodosa* rhizomes associated with clams were collected. *Codakia orbicularis* specimens for DMSP content across tissues were sampled in August 2021 in Guadeloupe (16.147778 N, −61.561111 W) by digging in a *Thalassia testudinum* seagrass bed. Clams were dissected into gills, visceral mass, foot, mantle, eggs, posterior and adductor muscle and frozen at −80°C until measurement.

### DMSP measurement

DMSP concentrations in clam tissues, sediment, and seagrass rhizomes were measured indirectly using the alkaline lysis method ^52^. Samples were treated with 300 μL of 10 M NaOH in 2 mL gas-tight vials, converting DMSP into DMS, and stored in the dark for 24 hours ^53^. DMS was quantified using a purge-and-trap gas chromatography system that used a flame photometric detector (Agilent 7890B GC) fitted with a 7693A autosampler and an HPINNOWax 30 m × 0.320 mm capillary column (Agilent Technologies J&W Scientific). The detection limit for DMS was 0.25 nmol. DMSP concentrations were calculated with an eightpoint calibration curve as in Curson *et al.,* ^53^ and normalised per gram of wet tissue or sediment. All DMSP concentration datasets failed the Shapiro-Wilk normality test (p < 0.05), so the Wilcoxon rank sum test was applied for statistical analyses. All statistical calculations were performed in R using the ggsignif package. The data was visualised using ggplot and tidyverse packages in R.

### 16S rRNA gene amplicon sequencing and analysis

The DNA for 16S rRNA gene amplicon sequencing was extracted from *Loripes orbiculatus* and *Loripinus fragilis* gills (9 samples each) obtained during environmental sampling and laboratory incubation (Supplementary methods, Sudo, 2023^54^) following the TRIzol™ (Thermo Fischer Scientific) extraction protocol for small quantities ^55^ with further modifications (Supplementary methods). Amplicons were generated and barcoded following a two-step PCR protocol ^56^. The first PCR step used the headed-primer set 341F and 785R ^57^, targeting the bacterial V3-V4 hypervariable region of the small subunit rRNA gene. Samples were sequenced at the Joint Microbiome Facility of the Medical University of Vienna and the University of Vienna (JMF). The resulting 16S rRNA gene amplicon reads were processed and analysed as in Pjevac *et al.*, 2021 ^56^. The 16S rRNA gene amplicon sequences were classified using the classifier implemented in DADA2 against the the SILVA database v.138.1 ^58,59^. The ASVs identified for the genera *Sedimenticola* and *Thiodiazotropha* were manually extracted and confirmed to be symbiotic ASVs via comparison to the available symbiotic 16S rRNA genes (metagenome-assembled genomes from Sudo *et al.*, 2024 ^55^).

### Isolation and cultivation of Ca. Endozoicomonas endolucinida

*Ca*. Endozoicomonas sp. was isolated from the gills of *L. orbiculatus* from Piran by grinding the gill tissue in filtered artificial seawater and subsequent serial dilution and spread plate method on marine broth 2216 (Difco) agar plates. White opaque colonies grew on agar plates in hypoxic conditions. The isolated species were identified by colony PCR and Sanger sequencing of the 16S rRNA gene of colonies in the third generation from isolation at Microsynth AG (Vienna, Austria) (Supplementary methods). The *Endozoicomonas* isolate was deposited in the DSMZ microbial collection under accession number XXXXX (currently awaiting confirmation from DSMZ).

### DMSP cleavage by Ca. Endozoicomonas endolucinida

To measure DMS production from DMSP by *Ca*. Endozoicomonas sp. isolates, each strain was first grown in Marine Broth (MB) media under anerobic conditions. All bacterial cultures were adjusted to an OD_600_ of 0.3 and the cells were inoculated at 20% (v/v) into 300 μL in 2 mL sealed glass vials with anaerobic-treated MB media in the presence and absence of 0.5 mM DMSP at 28 °C for 3 to 5 days. The production rate of DMS from DMSP was calculated based on incubations in 0.1mM DMSP as the sole carbon source in MBM media. DMS production in the headspace of vials was quantified by gas chromatography, as described above. The protein content in the cells was estimated by a Bradford method (BioRad) and the DMS production were expressed as pmol·mg^−1^ protein.

### Genome sequencing and assembly of Ca. Endozoicomonas endolucinida

High molecular weight DNA was extracted from *Ca*. Endozoicomonas endolucinida pure cultures using the Monarch HMW DNA extraction kit for tissue (NEB), with the specific protocol for Gram negative bacteria. Extracts were left at 4°C for at least one week before quantification and fragment estimation. Samples were diluted and equimolarly barcoded using the SQK-RBK114.96 (Oxford Nanopore Technologies) with the following protocol modifications: we increased the sample input to 80ng/sample, added 0.2 μL barcode/sample, and added 0.5 μL of rapid adaptor to the barcoded library. About 20fmol of a >60Kb library was loaded on a R10.4.1 flowcell (FLO-PRO114, Oxford Nanopore Technologies) and sequenced for 48 h on a Promethion P2 solo (Oxford Nanopore Technologies) using Minknow (v. 23.11.4, Oxford Nanopore Technologies). Flowcell light shields were used. Reads were basecalled using Dorado (v. 7.8.2) with super accuracy mode (r1041_e82_400bps_sup_v4.2.0). The nanopore raw reads were length and quality filtered using chopper (v. 0.6) (minlen=500bp, mean read score > Q15), and subsequently assembled using flye (v. 2.9.1, ^60^) with “–nano-hq” and polished once with medaka (v. 1.11.1, github.com/nanoporetech/medaka). Contigs <1000bp were removed. To compare the MAG from *Endozoicomonas* isolates to *Endozoicomonas* naturally occurring in lucinid clams, we assembled a new *Endozoicomonas* MAG from previously published Illumina sequencing data from *L. orbiculatus* in Piran under SRA accession number SRX26735717 ^61^, accordingly to Osvatic *et al.*, 2021 ^62^. The quality of assemblies was checked using CheckM (v. 1.2.3, ^63^), and classified using GTDBtk (v. 2.4.0.,^64^).

### Phylogenetic classification of Ca. Endozoicomonas endolucinida

For the phylogenetic analysis of the newly obtained symbiotic MAGs, we used additional publicly available MAGs from other *Endozoicomonas* species (Supplementary table SM1). All available MAGs underwent quality checks using CheckM’s taxonomy-specific workflow, with a gammaproteobacterial set of marker genes (v. 1.2.3, ^63^). Taxonomic classification was done using the GTDB-Tk v0.3.3 classification workflow. This workflow produced a concatenated amino acid sequence alignment of 120 highly conserved single-copy bacterial marker genes that were used for phylogenomic analyses ^64^. The phylogenetic tree was calculated using IQTree multicore version 2.1.2 ^65^, applying the default settings: auto substitution model detection - LG+I+G4 and 1000 ultra-rapid bootstraps (UFBoot) ^66^. The tree was rooted at *Marinobacter nauticus* and visualized using TreeViewer v. 2.2.0 ^67^. The resulting phylogenetic tree was used alongside Average Nucleotide Identity (ANI) analyses to identify species-level clades. FastANI v1.3 was employed to calculate the average nucleotide identity (ANI) between MAGs^68^, with a program-suggested 95% one-way ANI threshold used for species delimitation.

### Annotation and analysis of putative genes involved in organosulfur metabolism

Putative sulphur transformation genes from the lucinid symbionts, *Sedimenticola*, *Ca*. Thiodiazotropha and *Endozoicomonas* genomes were identified using HMSS2 ^69^ run on default settings, hmmsearch with Ddd hmm profiles from the HMSS2 tool with a higher e-value of 10^−10^, and using tblastn (BLAST+ 2.14.0) ^70,71^ search using a set of proteins described for organosulfur cycling, including DddP, DddW, DdhA, DmdA, DmdB, DmdC, DmdD, DmoAB, DmsA, DmsB, DmsC, MddA, MegL, MetB, MetH, Mtox, MddA, AcuH, Tmm, from UniProt and NCBI databases as a query with a maximal e-value 10^−20^ (Supplementary table SM2). The lucinid *Endozoicomonas* MAGs were annotated with the Rapid Annotation using Subsystem Technology (RAST) web server ^72^ using the RASTtk pipeline ^73^. The genes of interest were identified by the internal BLAST function in SEED ^74^. The gene models surrounding the *mtoX* locus were further verified using searches of the BLASTp against non-redundant protein sequences (nr) with an e-value cut-off of 10^−10^ ^75^ and InterProScan ^76^. The gene visualization was based on the annotation overview in the SEED genome browser and illustrated using Inkscape 1.4.

### Phylogenetic classification of DMSP synthesis candidate genes

For the search for candidate DSYB genes, we used *de novo* assembled transcriptomes of *Loripes orbiculatus* and *Loripinus fragilis* using transcriptomic data cleaned of symbiont reads from freshly collected clams in Piran, Slovenia as described in Sudo, 2023 (Supplementary methods)^54^. We searched for homologs of known DMSP synthesis proteins (DsyB/DSYB, DSYE, DsyGD/DsyG, TpMMT, MmtN and MMT) using BLAST using blastp (BLAST+ 2.14.0; ^75^ (Supplementary table SM2). We also run the hmm search tool in HMMER v3.3.2 ^77^ with an e-value cutoff of 10^−10^, and DsyB/DSYB and DsyGD/DSYE hmm profiles obtained from Curson *et al.*, 2018 and Wang *et al.*, 2024 ^12,78^. The alignment was constructed in Clustal Omega v. 1.2.4 ^79^ using default settings together with sequences representing other members of related methyltransferase families, phosphatidylethanolamine N-methyltransferase (pmt), acetylserotonin O-methyltransferase (ASMT), and dTTP/UTP pyrophosphatase (maf), as well as representatives of homologs of known DMSP synthesis proteins as mentioned above (Supplementary table SM2). The tree was calculated using IQ-Tree multicore version 2.1.2 ^65^ using the default settings (auto substitution model detection and 1000 ultra-rapid bootstraps (UFBoot) ^66^. The tree was rooted at the midpoint and visualised using TreeViewer v. 2.2.0 ^67^.

### Live incubations of Loripes orbiculatus

To investigate the impact of microbial DMSP degradation and anoxia stress, we incubated *L. orbiculatus* holobionts in triplicate in oxic and anoxic conditions for 48 h in the dark with and without exposure to an antibiotic cocktail (Supplementary figure SM1, Supplementary methods).

## Results and discussion

### Some lucinids are ‘hotspots’ of DMSP

To explore the role of DMSP in lucinid holobionts, we quantified DMSP in two co-occurring clam species (n=107) from Piran, Slovenia, and their surrounding habitats (Figure 1). DMSP concentrations in *Loripes orbiculatus,* reaching up to 175.46 nmol g^−1^, were consistently higher (p ≤ 0.001) than those in the environment (p ≤ 0.01). In contrast, DMSP concentrations in *Loripinus fragilis* were not significantly different in than in the surrounding environment (Figure 1). Differences between lucinid species may result from varying rates of host accumulation through diet, microbial/host species-dependent DMSP production within the holobiont and/or differing DMSP catabolic rates in the different hosts. High DMSP concentrations similar to those measured in *L. orbiculatus* have been observed in some species of marine animals, and are also attributed to dietary intake or production by phototrophic symbionts (Supplementary figure SR1A) ^9^.

**Figure 1:**
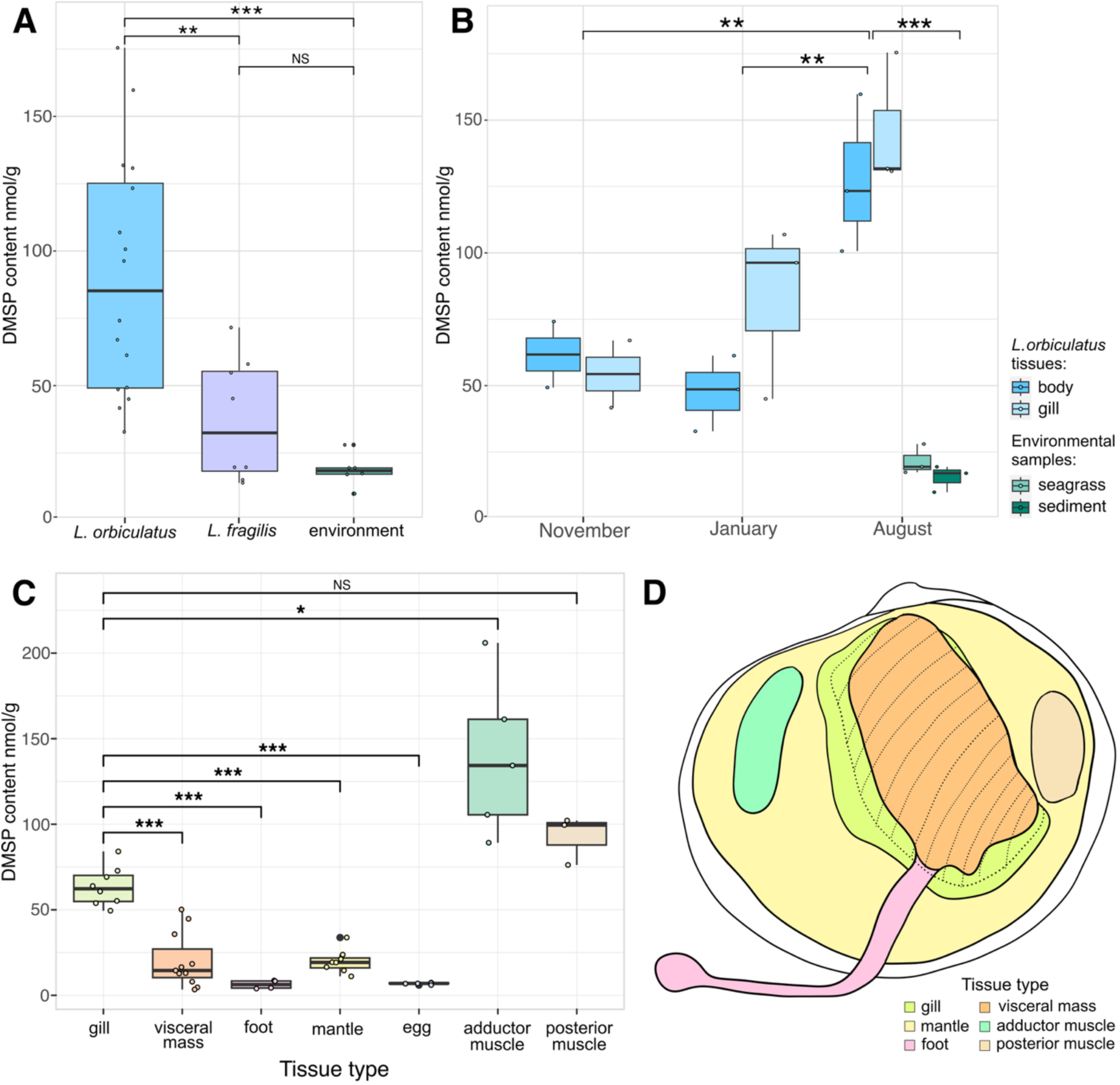
DMSP concentrations in (A) all samples from *Loripes orbiculatus*, *Loripinus fragilis*, and environmental samples (seagrass roots and sediment), (B) *Loripes orbiculatus* tissues and environmental samples over sampling months, (C) various tissues of *Codakia orbicularis* from Guadeloupe. (D) Schematic illustration of lucinid tissues investigated in this study, the colouring reflects the tissue labelling in panel C. The main population of symbionts is in the gills. In this image, one gill is shown overlapping with the visceral mass, marked here as dashed lines. Significant differences are labelled as “*” for p ≤ 0.05, “**” for p ≤ 0.01, and “***” for p ≤ 0.001.

### The seasonality of DMSP accumulation in lucinids

To investigate the source of DMSP in lucinids, we measured concentrations in the clam tissues and their seagrass and sediment environment across seasons. In filter-feeding mussels such as *Mytilus edulis*, which lack chemosynthetic symbionts, DMSP levels peak in summer ^80^. Similarly in *L. orbiculatus*, DMSP levels varied seasonally (Figure 1), with significantly higher (p ≤ 0.01) concentrations in summer (August) compared to autumn and winter (November and January). There were no significant differences between the gill and body tissues. However, lucinids derive only 20–39% of their nutrition from filter feeding ^81^, and the known seasonal variation in *L. orbiculatus’s* reliance on filter feeding shows the opposite trend to what we would expect if the DMSP were obtained from filter-feeding. *L. orbiculatus* filter-feeding peaks in early autumn and winter, while DMSP concentrations were highest in summer (Figure 1) ^81^. Furthermore, in August, DMSP concentrations in Piran sediment and seagrass were up to 18 times lower (p ≤ 0.001) than in *L. orbiculatus* tissues. Given these observations, we hypothesize that the high DMSP concentration in *L. orbiculatus* was due to internal synthesis and not from its food.

### DMSP in lucinid holobionts is independent of symbiont abundance and unlikely of bacterial origin

Recent studies demonstrated that some marine heterotrophic bacteria can synthesise DMSP ^50,52^, thus, we investigated whether bacterial symbionts could be the source of DMSP in lucinids. Although symbionts could be widespread in the body of lucinids, the gills host the majority of population of symbionts^82^. We therefore quantified DMSP across different tissues to see if DMSP concentrations were highest in the gills, which could indicate synthesis by the symbionts. In both *Loripes orbiculatus* and *Loripinus fragilis*, DMSP concentrations were consistent across tissues despite major differences in symbiont abundances, indicating that the sulphur-oxidizing symbionts were unlikely responsible for DMSP production in these clams (Figure 1A, Supplementary figure SR1B). *L. orbiculatus* and *L. fragilis* are typically quite small, with shell lengths rarely more than 2 cm. It was therefore impossible to dissect and collect enough tissue to measure DMSP concentrations in all body parts. To gain a more comprehensive understanding and pinpoint the possible origin of DMSP in lucinid hosts, we measured DMSP in all body parts of *Codakia orbicularis*, a typically larger lucinid species. DMSP concentrations were significantly higher in the symbiont-harbouring gills compared with tissues not known to host a large population of symbionts, including the visceral mass (p ≤ 0.001), foot (p ≤ 0.01), mantle (p ≤ 0.001), and eggs (p ≤ 0.01). However, DMSP levels were overall highest and significantly higher than the gills in the adductor muscle (p ≤ 0.01) and posterior muscle (p ≤ 0.05), which are considered symbiont-free. Similar observations were reported for non-symbiotic abalone *Haliotis midae*, where the highest DMSP concentration was measured in the adductor muscle ^83^.

Furthermore, using BLAST and HMM-based searches, we could not find any candidate genes for DMSP synthesis in the genomes of the major lucinid symbionts, *Ca.* T. luna, *Ca.* S. endoloripinus, or the newly isolated *Ca*. Endozoicomonas endolucinida (discussed below). Based on the absence of DMSP synthesis genes in lucinid symbiont genomes, no detectable DMSP production by the *Ca*. E. endolucinida isolate (Supplementary results and discussion), as well as the lack of a symbiont-related pattern in DMSP concentrations within host tissues, we propose that the symbionts in lucinid holobionts were unlikely to be the DMSP source. Instead, it was more plausible that the lucinid host was responsible for DMSP production. Indeed, *Loripes orbiculatus* accumulated significantly more DMSP when incubated in anoxic seawater with antibiotics and no external food source compared to controls, supporting the clam as the DMSP source, rather than its food or associated microbiome (Figure 2, Supplementary results and discussion). Both, anoxia as well as addition of antibiotics, either by state of dysbiosis or by direct stress caused by antibiotics, may have triggered increased DMSP production by the host.

**Figure 2:**
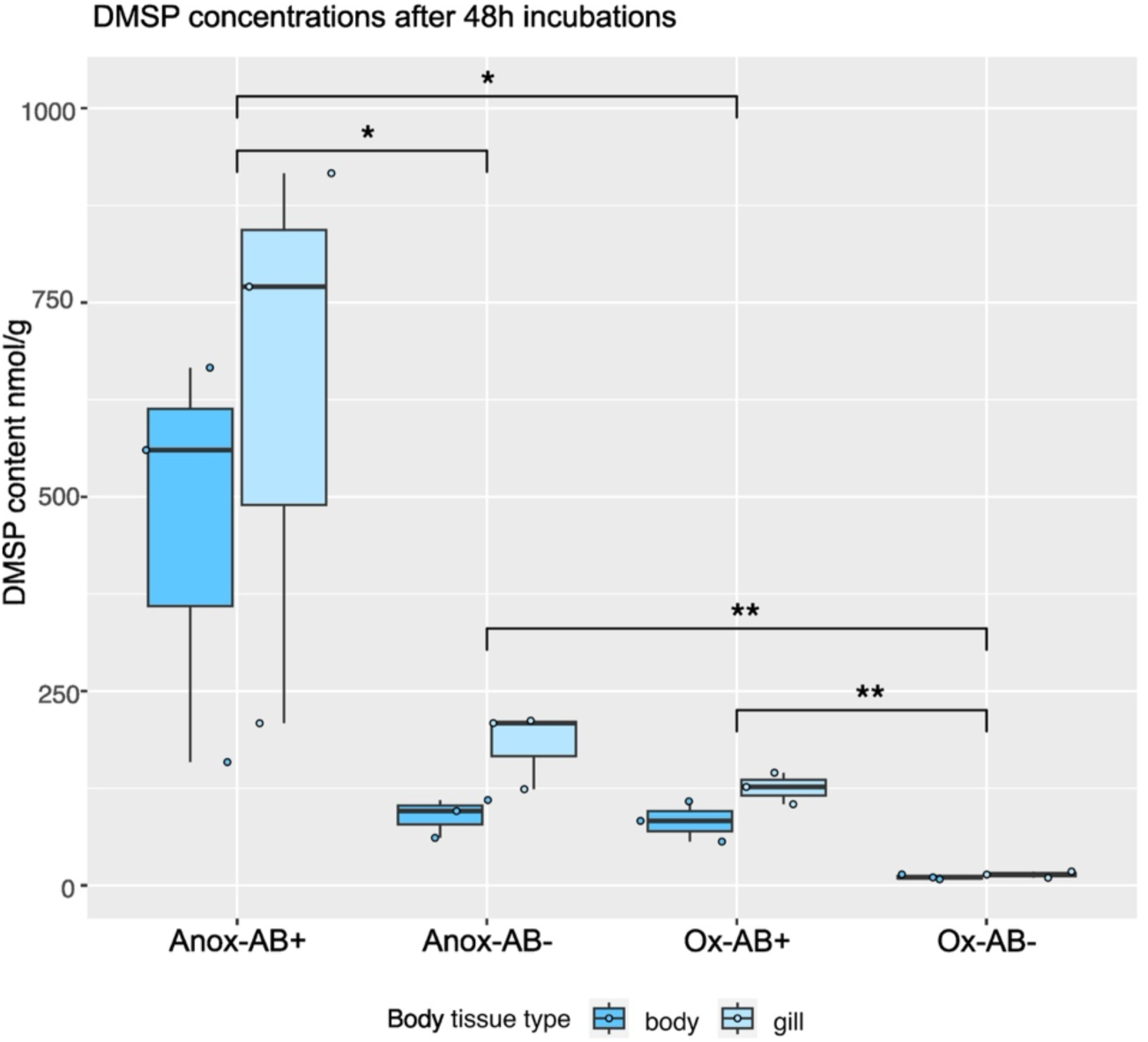
DMSP content in *Loripes orbiculatus* tissues in the whole holobiont antibiotic treatment experiment. The treatments included anoxic incubation with antibiotics (Anox-AB+) or without them (Anox-AB-) and oxic incubations with antibiotics (Ox-AB+) or without them (Ox-AB-). The significance is marked as “*” for p ≤ 0.05, “**” for p ≤ 0.01.

### Potential mechanisms of DMSP synthesis in lucinid hosts

DMSP synthesis has been identified in diverse taxa, including algae, numerous bacteria, and the saltmarsh grass *Spartina* ^6,84^. Evidence for DMSP production in animals, such as *Acropora* corals, has emerged only recently, and the underlying mechanisms are not yet characterized in detail. In these corals, methionine-derived DMSP is proposed to be synthesized by DSYB enzyme, an MTHB S-methyltransferase^13^. To identify potential DMSP synthesis genes in lucinid clams, we searched *de novo* assembled transcriptomes of specimens collected in their natural habitat in Piran for homologs of known DMSP synthesis genes (DsyB, DSYB, DSYE, DsyG, DsyGD, TpMMT, MmtN and MMT). Significant hits with a cutoff of 10^−10^ were detected in both lucinid species, *Loripes orbiculatus* and *Loripinus fragilis*, though with low amino acid identity: DN147177 (27.0 - 27.5%) and DN86992 (23.4 - 28.2%) in *L. orbiculatus*, and DN14881 (20.6 - 28.1%) and DN3985 (23.4 - 33.9%) in *L. fragilis*. Phylogenetic analyses placed DN147177 and DN14881 sequences in a sister clade distinct from canonical DSYB proteins, yet the most significant hits include DsyB homologs, and their domain structures closely resemble DSYB enzyme (Figure 3, Supplementary Figures SR5–SR6). Given that experimentally verified DSYB enzymes can share as little as ∼33% amino acid identity, functional annotation within this enzyme family remains challenging ^12,52^ and many pathways and enzymes likely remain to be discovered. We attempted functional characterization of these candidate proteins, but results were inconclusive. Despite this, these sequences represent promising candidates for further investigation of animal-associated DMSP synthesis.

**Figure 3.**
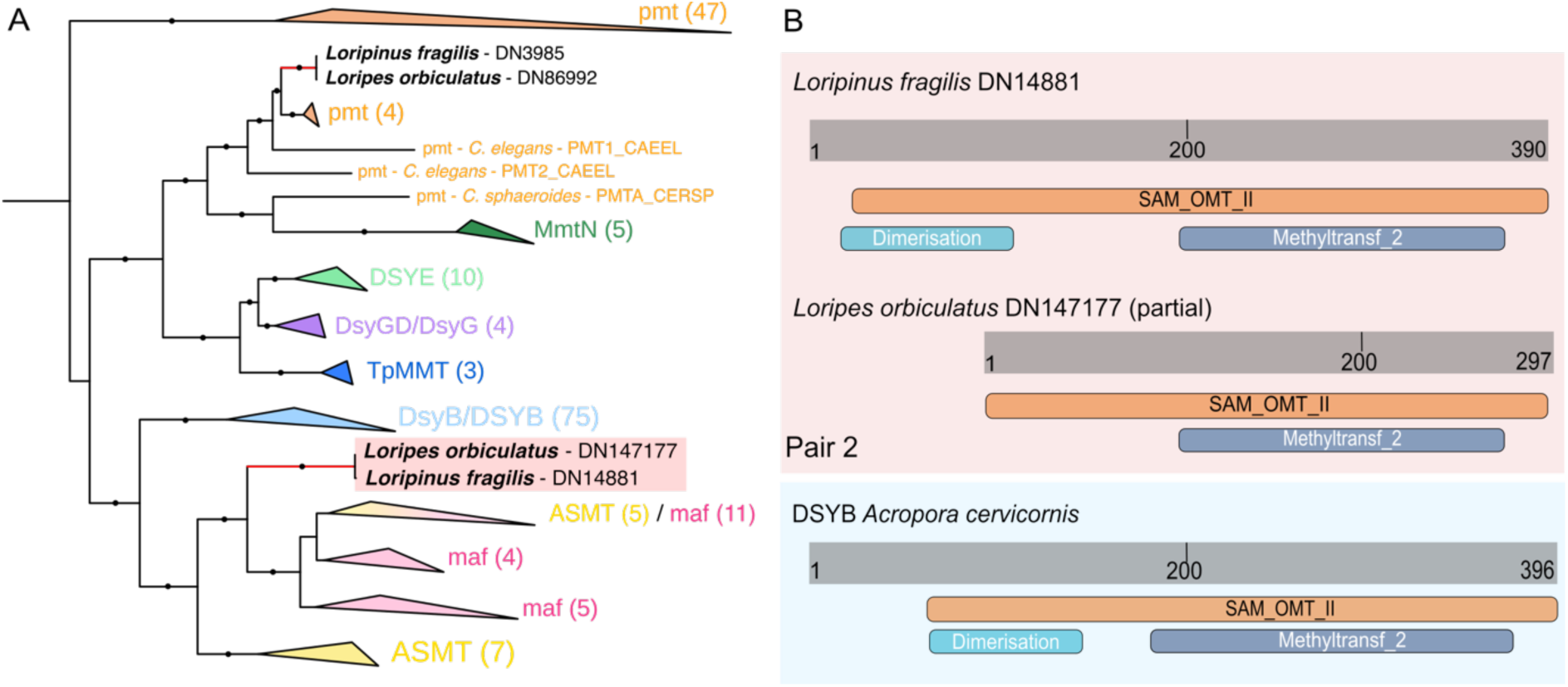
(A) A maximum likelihood phylogenetic tree of candidate DSYB and related gene families reconstructed using the best fit model Q.pfam+I+G4. Black circles indicate branch support over 85%. Proteins in this tree are methylthiohydroxybutryate methyltransferases (DSYB/Dsyb, blue; DsyGD/DsyG, violet; DSYE, light green; TpMMT, dark blue, MmtN, dark green), lucinid putative DSYB (black), Phosphatidylethanolamine N-methyltransferase (pmt, orange), Acetylserotonin Omethyltransferase (ASMT, yellow), and dTTP/UTP pyrophosphatase (maf, pink). Numbers in brackets indicate the number of sequences in the collapse clade. (B) Protein domains found by InterProScan for the pair 2 candidate DSYBs in *Loripes orbiculatus* and *Loripinus fragilis* clams resembles the domains in DSYB described in coral *Acropora cervicornis,* i.e. the O-methyltransferase domain, also known as Methyltransf_2 (PF00891 or IPR001077) as well as O-methyltransferase dimerisation domain (PF08100 or IPR012967) assigning this peptide as SAM-dependent O-methyltransferase class II-type profile (PS51683 or IPR016461).

### Ca. Endozoicomonas endolucinida is a consistent member of the Loripes orbiculatus microbiome

The microbiome of lucinid clams, though dominated by the sulphur-oxidizing symbionts, has been shown to include various other ‘secondary’ symbionts, usually of much lower relative abundance ^35,51,82,85^. To identify microbes potentially involved in organosulfur cycling, we analysed the diversity of *L. orbiculatus* and *L. fragilis* gill microbiomes by 16S rRNA gene amplicon sequencing. The microbiomes were dominated by the previously characterized sulphur-oxidizing symbionts *Ca.* T. luna and *Ca.* S. endoloripinus, respectively (Figure 4). *Spirochaeta* and *Endozoicomonas* were prominent in both clam species, while *Shewanella* was only present in *L. orbiculatus* gills. Additionally, *L. orbiculatus* harboured low-abundance bacteria from *Flavobacteriaceae, Izemoplasmatales*, *Margulisbacteria*, and *Rhizobiales,* which were not detected in *L. fragilis*. *Endozoicomonas* made up a larger proportion of sequence reads in *Loripes* (up to 39.3% of sequences, averaging 19%, st. dev. 10%) compared to *Loripinus* (average of 1.5%, st. dev. 1.3%). The presence of *Endocoizomonas* ASVs in lucinid gill microbiomes from our study was consistent with previous reports of these bacteria being enriched in the gill microbiomes of other lucinid species, including *Ctena orbiculata*, *Anodontia alba*, *Euanodontia ovum*, *Lucinisca nassula*, and *Codakia orbicularis* ^35,51,85^. *Endocoizomonas* were also seen in other sulphur-oxidising symbioses such as *Thyasira cf. gouldi* and *Alvinoconca sp*.^86^, implying their possible conserved role in chemosymbiotic associations. With *Endozoicomonas* being associated with higher levels of DMSP we proposed that they might either synthesize and or cycle DMSP. To investigate this potential, we isolated and sequenced the genome of an *Endozoicomonas* strain from *L. orbiculatus*. With long-read sequencing, we were able to produce a high-quality genome on four contigs with 99.44% completeness and 1.69% contamination, to our knowledge, the best quality genome of a bivalve-associated *Endozoicomonas* obtained so far (Supplementary table SR3). Phylogenetic analysis confirmed that the isolated *Endozoicomonas* strain was identical to the symbiont present in *L. orbiculatus* gills, as all five 16S rRNA gene sequences from the draft genome of the isolate clustered with the five ASVs recovered directly from *L. orbiculatus* gill tissue (Supplementary figure SR3). Furthermore, our isolate clustered as a distinct clade in both 16S rRNA (Supplementary figure SR3) and genome-based phylogenetic trees (Figure 5), indicating that this isolate likely represented a novel *Endozoicomonas* species. This was further supported by ANI analysis (Supplementary table SR4), with other *Endozoicomonas* genomes sharing at most 78.38% similarity to our isolate. We therefore propose the species name *Ca*. Endozoicomonas endolucinida, highlighting the origin of our isolate from a lucinid clam.

**Figure 4:**
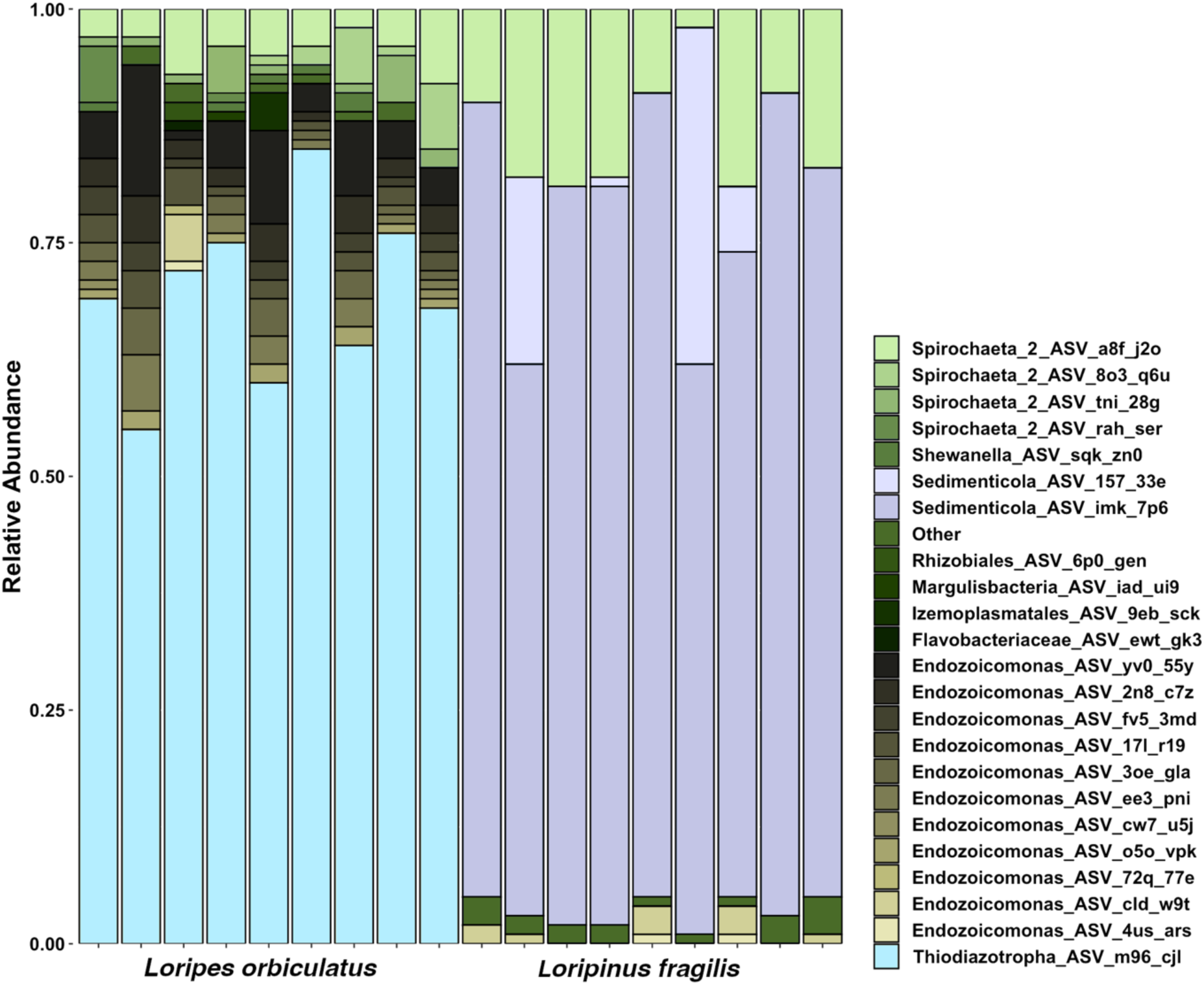
Relative abundance of 16S rRNA gene ASVs found in *Loripes orbiculatus* and *Loripinus fragilis* gills. Each unique ASV is shown in a different colour. The sulphur-oxidizing symbionts are *Ca.* T. luna (labelled as Thiodiazotropha_ASV_m96_cji, blue) and *Ca.* S. endoloripinus (Sedimenticola_ASV_157_33e and Sedimenticola_ASV_imk_7p6, violet).

**Figure 5:**
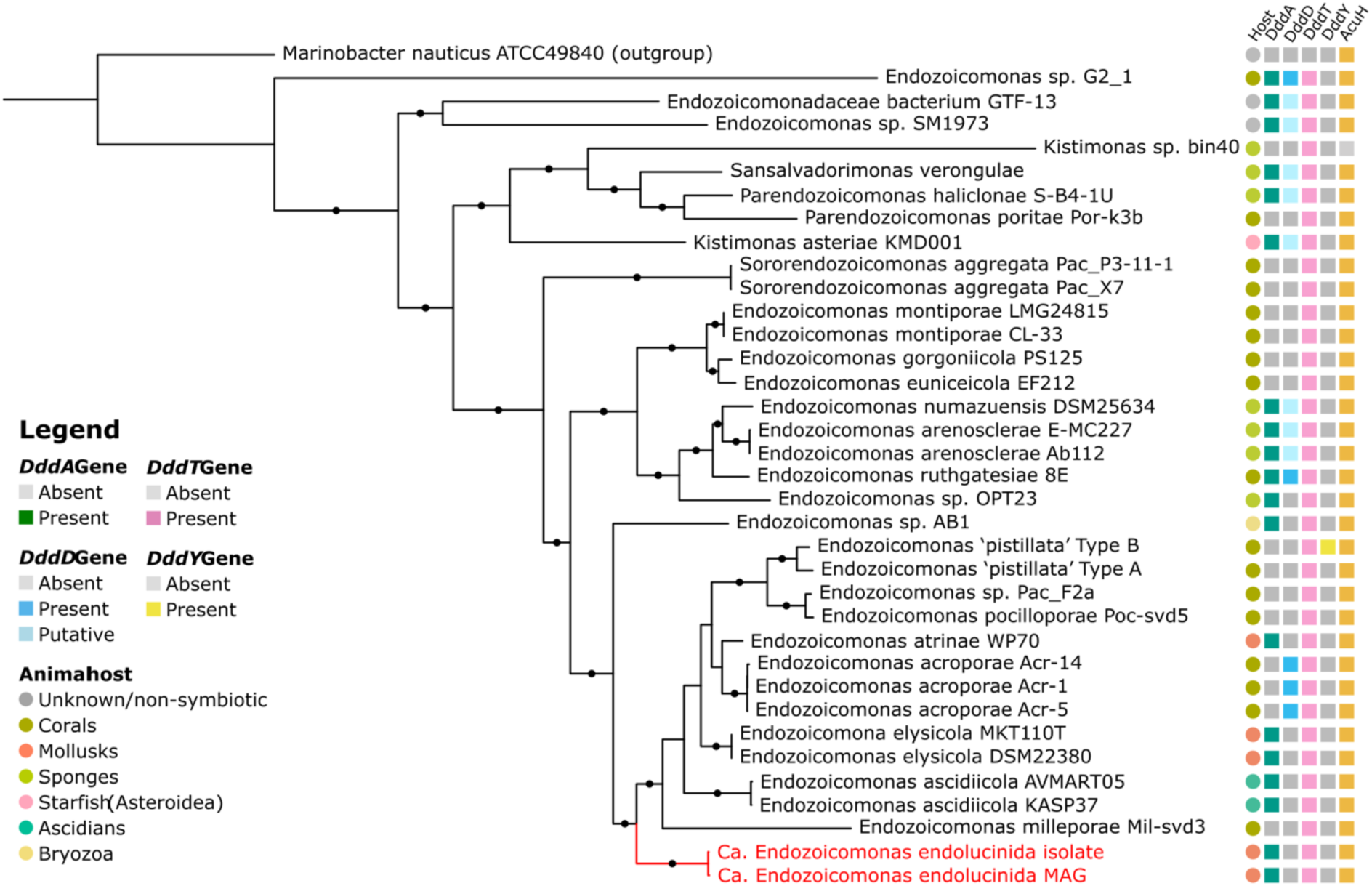
A maximum likelihood phylogenetic tree was reconstructed from GTDB’s multisequence alignment using the best-fit model LG+I+G4. Circles indicate bootstrap support values above 95%. Squares indicate the presence of DMSP catabolism-related genes, the host group is indicated by coloured circles according to the colour legend. The tree is rooted at *Marinobacter nauticus*.

### Endozoicomonas catabolizes DMSP via an as-yet-unknown lysis pathway in the lucinid holobiont

The sister clade to *Ca*. E. endolucinida contains *Endozoicomonas* species found in other mollusks, ascidians, and corals, some of which have *dddD or dddY* genes for DMSP lysis (Figure 5, ^87,88^). Surprisingly, though no primary DMSP cleavage or demethylation genes were detected in the high-quality *Ca*. E. endolucinida genome despite it containing homologous genes for a DddT DMSP transporter ^89,90^ and DddA which is an alcohol dehydrogenase involved in the processing of the 3-C DMSP cleavage intermediate 3-OH-propionate to malonate semialdehyde ^91^. Furthermore, *Ca*. E. endolucinida encoded an AcuH enoyl-CoA hydratase family, which is required for detoxification of acrylate produced during DMSP lysis (Supplementary table SR2, ^25^). Consistent with these bioinformatic predictions, we confirmed that *Ca*. E. endolucinida could import DMSPd and concentrate it to estimated 0.90-3.40 μM cellular levels (Supplementary results and discussion). Furthermore, all *Ca*. E. endolucinida isolates cleaved DMSP to DMS and acrylate, as indicated by detection of DMS after 86h of incubation (28.9-123.5 nmol mg⁻¹ protein) and specific production rates of 5.59-23.93 nmol mg⁻¹ protein min⁻¹ in Marine Broth media and 12.24 to 92.22 pmol mg⁻¹ min⁻¹ with DMSP as a sole carbon source compared with no detectable DMS production in the media controls or in isolates incubated without DMSP (Figure 6). These activities fall within the range reported for known bacterial DMSP lyases, although the apparent rates may be underestimated due to a longer incubation period and non-saturating substrate concentrations ^92^. Given the significant biodiversity of characterized DMSP-cleaving enzymes, and the observation that several taxa (e.g. Actinobacteria) exhibit DMSP lyase activity despite lacking canonical *ddd* genes, we hypothesized that *Ca*. E. endolucinida contains a novel, undescribed DMSP lyase ^93^. These observations refine the hypothesis by Pogoreutz *et al.*, 2022 ^88^, who proposed that while DMSP degradation may be an important metabolic trait in marine bacterial symbioses, it is not a universal feature among *Endozoicomonas*. Our findings suggest that DMSP cleavage capacity may extend beyond the previously characterized *Endozoicomonas* lineages, indicating that this metabolic trait could be more broadly distributed within the Endozoicomonadaceae.

**Figure 6:**
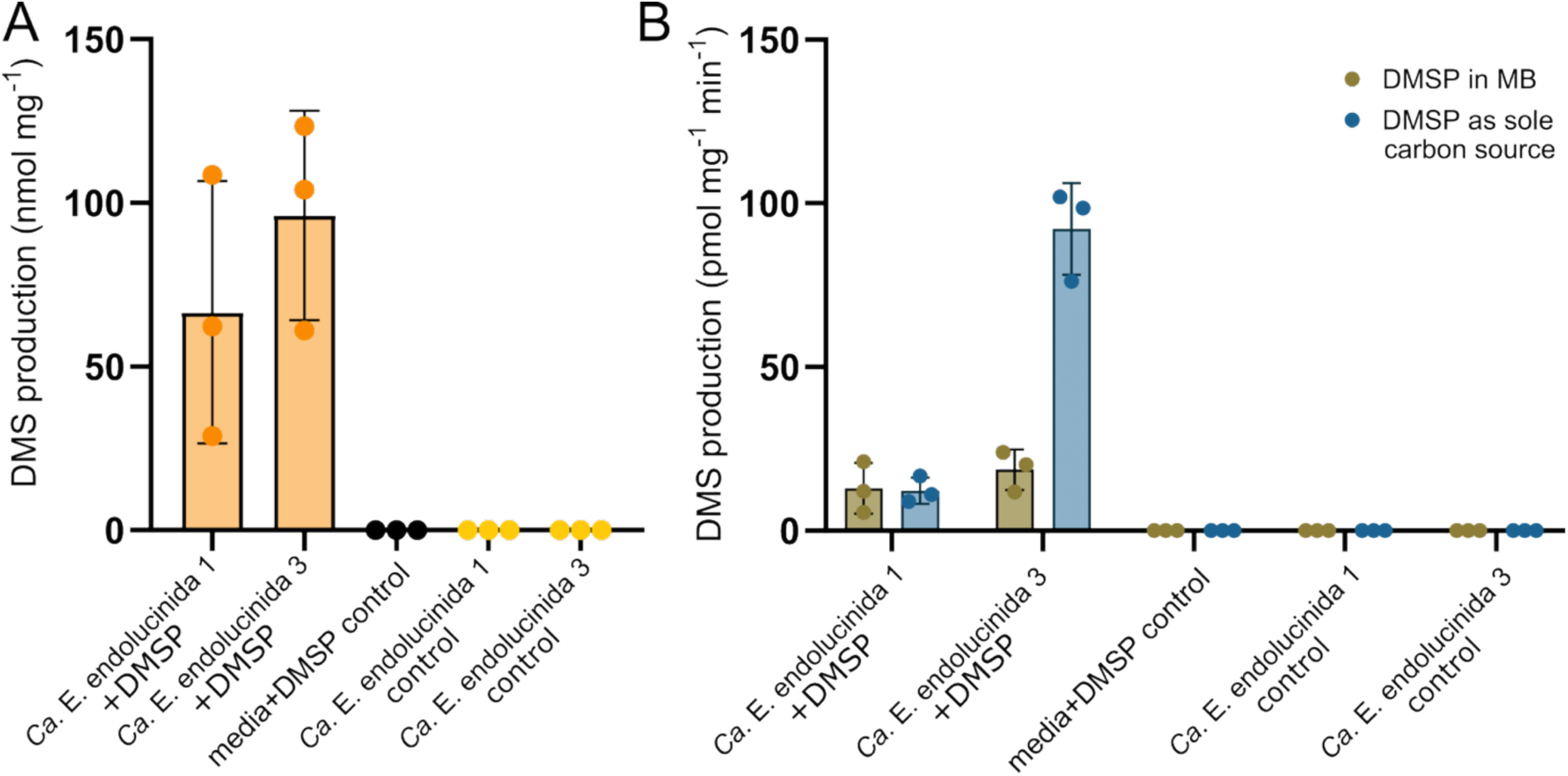
DMSP cleavage by *Ca*. E. endolucinida. (A) Total DMS production during incubations in Marine Broth (MB) media. Both clonal isolates 1 and 3 (orange) produced DMS after incubation with addition of DMSP, while no addition of DMSP (yellow) and control samples with media only and addition of DMSP (black) show no DMS production. (B) DMS production rates by *Ca*. E. endolucinida in MB media (brown) and minimal media with DMSP as sole carbon source (blue).

*Lucinid sulphur-oxidizing symbionts may link inorganic and organosulfur transformations*.

Inorganic sulphur oxidation is the basis of symbiotic interaction in lucinid holobionts ^44^. However, the role they or any other chemosynthetic symbiosis may play in marine organosulfur cycling has not been considered in detail, despite DMSO reductase being previously reported as one of the highest expressed genes (Supplementary table SR2)^48^ and suggested as a potential alternative electron sink. We found homologs of the *dmsABC* gene complex in symbionts of *L. orbiculatus* and *L. fragilis,* implicating this chemosynthetic symbiosis in coastal organosulfur cycling. The gaseous DMS, produced through DMSO reduction by the sulphur-oxidising symbionts or by potential lysis of DMSP by *Ca*. E. endolucinida could be released into the seawater and contribute to the marine DMS emissions. DMS could also be potentially reoxidized back to DMSO as an intermediate step of multicomponent DMS monooxygenase detected in the sulphur-oxidizing symbiont genomes of both lucinid species (Supplemental results and discussion).

Alternatively, DMS produced by the symbionts could be further catabolised by *Ca. T. luna*, whose genome encodes a candidate MtoX methanethiol oxidase (MTO) with 60% amino acid identity to these characterized enzymes. MTO activity is common in bacteria utilising MeSH, but also on DMS, as a sole carbon and energy source ^94,95^. As with bona fide *mtoX* genes, *Ca*. T. luna *mtoX* was predicted to be in an operon with SCO1/SenC and MauG-encoding genes (Figure 7) that are essential in the regulation of *mtoX* expression, MTO maturation, and posttranslational modification ^94^. The presence of this gene cluster implied that *Ca*. T. luna may possess the enzymatic machinery to oxidize DMS-derived MeSH to formaldehyde and hydrogen sulphide, as described in other methylated sulphur oxidizers such as *Hyphomicrobium denitrificans* and *Ruegeria pomeroyi* ^94,96^. Note, the enzyme responsible for DMS demethylation is currently unknown in bacteria with MTO. Both, sulphide and formaldehyde can be further detoxified and oxidized via the complete tetrahydromethanopterin-dependent pathway and various sulphur oxidation pathways, all encoded in the *Ca*. T. luna genome (Figure 8, Supplementary figure SR7, ^55^). Since MeSH could be toxic to animal host, this metabolic capacity of MeSH oxidation could be not only beneficial for the symbiont but also the host. The source of MeSH in this symbiotic system could be (1) methylation of H_2_S by other microbes in their environment, (2) methionine catabolism by methionine γ-lyase *MegL,* or (3) mitochondrial degradation of methionine by the host, which in mammals regulates excess methionine ^97^. All analysed symbionts encode *megL* in their genomes (with 43.71% aa identity in *Ca*. S. endoloripinus, 42.97% in *Ca*. T. luna and 43.04% in *Ca*. E. endolucinida compared to *megL* from *Pseudomonas putida*), which is common to most bacteria. There is no direct evidence of the role of methionine in lucinid symbioses, however, accumulation of cellular L-Met is potentially toxic ^98^. The symbiotic MegL could alleviate this toxicity by breaking down excess L-Met, providing an additional pathway for detoxifying or recycling reduced sulphur compounds within the holobiont and thereby expanding its known metabolic repertoire.

**Figure 7:**
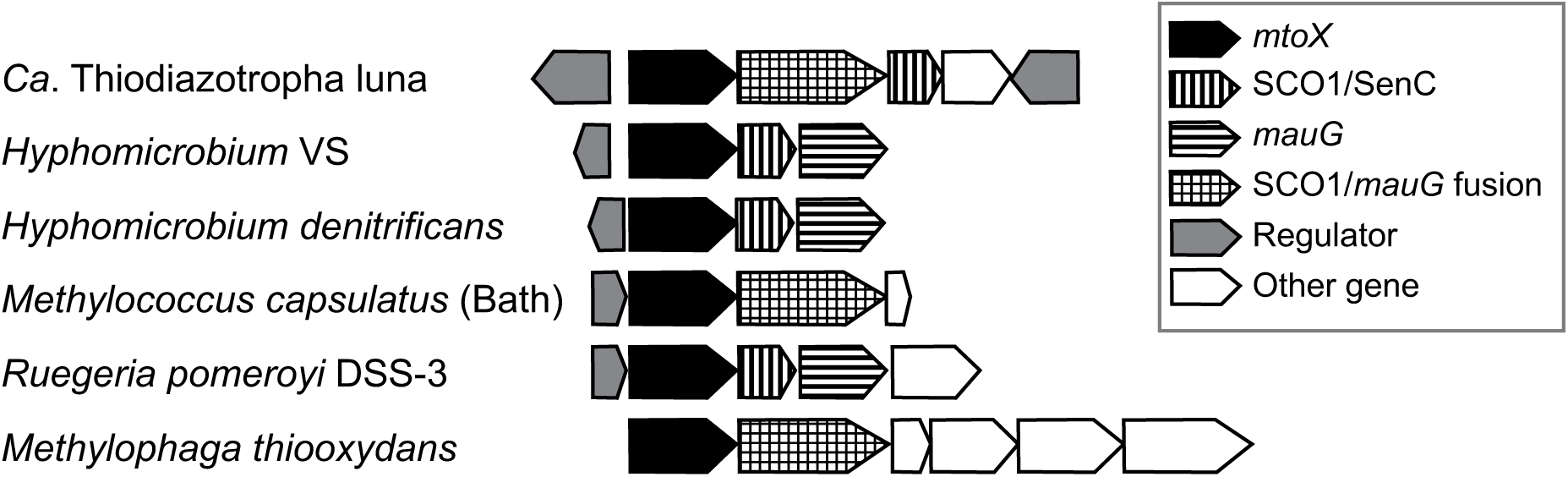
Genomic context of the *mtoX* gene in *Ca*. Thiodiazotropha luna and known MeSH metabolizing bacteria. Just as in *Methylococcus* and *Methylophaga*, *Ca*. Thiodiazotroha luna encodes a fusion protein with SCO1/SenC and mauG domains. After Eyice *et al.*, 2018 ^94^.

**Figure 8:**
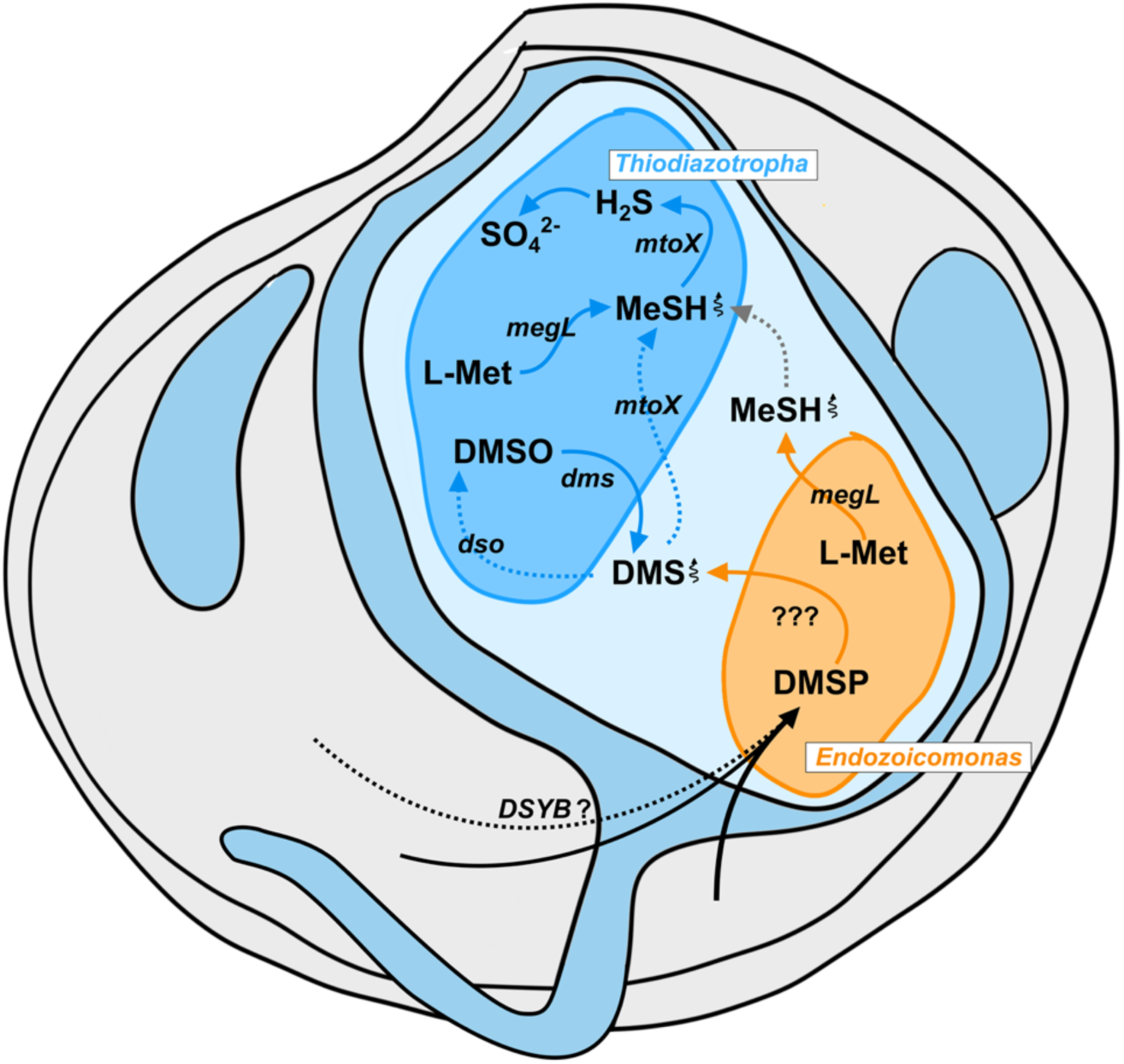
Proposed symbiotic organosulfur cycling in *Loripes orbiculatus* and possibly other lucinid clam species. Coloured arrows represent putative pathways in the sulphur oxidizer *Ca.* T. luna (blue), *Ca*. E. endolucinida (orange) and *L. orbiculatus* host (black). Pathways of lower confidence are marked by dashed lines. *Ca.* T. luna (blue), and *Ca*. E. endolucinida (orange) were both detected in the gills of *L. orbiculatus*, but the exact location of *Ca*. E. endolucinida is not yet known.

### Functional partitioning between Thiodiazotropha and Endozoicomonas may facilitate catabolism of host-derived DMSP

Considering the ability of *Ca. E. endolucinida* to catabolize DMSP to DMS (Figure 6), the presence of multiple pathways for metabolizing DMS, MeSH, DMSO, and methionine in the sulphur-oxidizing *Ca. T. luna* (Figure 8), and the consistent detection of multiple symbiotic partners in all individuals examined (Figure 4), the complete DMSP degradation may be possible within the *Loripes orbiculatus* holobiont but it would require coordinated catabolism of multiple partners. Such functional partitioning is well known from insect symbioses, where processes such as vitamin and amino acid biosynthesis depend on a “community” of complementary symbionts (e.g. *Buchnera* and *Serratia* in aphids ^99^, *Sulcia* and *Baumannia* or *Hodgkinia* in cicadas ^100^). Collaborative bacterial DMSP degradation in *L. orbiculatus* (Figure 8) is consistent with experimental results from live incubations of the holobiont, in which addition of antibiotics in both oxic and anoxic conditions resulted in significantly higher DMSP concentrations (Figure 2, Supplementary results and discussion). We propose that lucinid clams may synthesize DMSP, especially under stress, a trait observed in only a few recently reported animal species. The novel *Endozoicomonas* species isolated here, and the well-studied sulphur-oxidizing symbionts may cooperate to degrade excess DMSP, potentially protecting the host from pathogenic bacteria. Lucinid holobionts may not only have an impact on organosulfur cycling in the sediments of their local Mediterranean or Caribbean habitats, but also at global scale given their worldwide distribution. Still, large-scale quantification and further studies are required to account for the variations observed across different lucinid species, as seen in this study. Additionally, given the diverse beneficial roles of *Endozoicomonas*, e.g., nutrition acquisition, antimicrobial activity, microbiome structuring and defence against pathogens, our novel isolate paves the way for the first culture-dependent studies of beneficial interactions within lucinid holobionts.

## Acknowledgements

Financial support was provided by the ERC Starting Grant EvoLucin (grant number 802494), a Vienna Research Grant for Young Investigators from the Vienna Science and Technology Fund (WWTF, VRG14-021), the Austrian Science Fund project MAINTAIN DOC 69 doc.fund., the Austrian Academy of Sciences, and the FWF Cluster of Excellence Microplanet. Work in JDT’s laboratory was additionally funded by NERC (NE/P012671, NE/S001352, NE/X000990 and NE/X014428) and the Leverhulme Trust (RPG-2020-413). The Joint Microbiome Facility (JMF) of the Medical University of Vienna and the University of Vienna (project ID JMF-2209-07) conducted most of the sequencing, with invaluable assistance in sample processing provided by Petra Pjevac and Gudrun Kohl. Computational resources essential to this study were provided by the Life Science Computer Cluster at the University of Vienna. We also thank the Marine Biological Station Piran, Slovenia, and the scuba diving team for their support during the fieldwork and laboratory incubations.

## Author contributions

Conceptualization - M.S., J.M.P., J.D.T.

Sample collection - M.S., O.G., J.M.P.

Data collection and analysis - M.S., C.S., O.C., O.Z., J.S., J.O.

Writing - M.S., J.M.P., with contributions from all co-authors

Supervision - J.M.P., J.D.T.

## Data Availability

The data (raw 16S rRNA gene sequences, the genome of *Ca*. E. endolucinida together with the respective reads libraries, RNAseq reads data) are deposited in the NCBI database under BioProject PRJNA1227762.

## References

1. Ksionzek, K. B. et al. Dissolved organic sulfur in the ocean: Biogeochemistry of a petagram inventory. Science 354, 456–459 (2016).

2. Sunda, W., Kieber, D. J., Kiene, R. P. & Huntsman, S. An antioxidant function for DMSP and DMS in marine algae. Nature 418, 317–320 (2002).

3. Kirst, G. O. Osmotic Adjustment in Phytoplankton and MacroAlgae. in Biological and Environmental Chemistry of DMSP and Related Sulfonium Compounds (eds Kiene, R. P., Visscher, P. T., Keller, M. D. & Kirst, G. O.) 121–129 (Springer US, Boston, MA, 1996).

4. DeBose, J. L., Lema, S. C. & Nevitt, G. A. Dimethylsulfoniopropionate as a Foraging Cue for Reef Fishes. Science 319, 1356–1356 (2008).

5. Seymour, J. R., Simó, R., Ahmed, T. & Stocker, R. Chemoattraction to Dimethylsulfoniopropionate Throughout the Marine Microbial Food Web. Science 329, 342–345 (2010).

6. Carrión, O. et al. Molecular discoveries in microbial DMSP synthesis. in Advances in Microbial Physiology vol. 83 59–116 (Elsevier, 2023).

7. Bates, T. S., Charlson, R. J. & Gammon, R. H. Evidence for the climatic role of marine biogenic sulphur. Nature 329, 319–321 (1987).

8. Raina, J.-B., Dinsdale, E. A., Willis, B. L. & Bourne, D. G. Do the organic sulfur compounds DMSP and DMS drive coral microbial associations? Trends in Microbiology 18, 101–108 (2010).

9. Van Alstyne, K. L., Dominique, V. J. & Muller-Parker, G. Is dimethylsulfoniopropionate (DMSP) produced by the symbionts or the host in an anemone–zooxanthella symbiosis? Coral Reefs 28, 167–176 (2009).

10. Hill, R. W., Dacey, J. W. H. & Edward, A. Dimethylsulfoniopropionate in giant clams (Tridacnidae). Biological Bulletin 199, 108–115 (2000).

11. Aguilar, C. et al. Transcriptomic analysis of the response of *Acropora millepora* to hypoosmotic stress provides insights into DMSP biosynthesis by corals. BMC Genomics 18, 612 (2017).

12. Curson, A. R. J. et al. DSYB catalyses the key step of dimethylsulfoniopropionate biosynthesis in many phytoplankton. Nat Microbiol 3, 430–439 (2018).

13. Raina, J.-B. et al. DMSP biosynthesis by an animal and its role in coral thermal stress response. Nature 502, 677–680 (2013).

14. Hill, R. W., Dacey, J. W. H. & Krupp, D. A. Dimethylsulfoniopropionate in Reef Corals. Bulletin of Marine Science 57, 489–494 (1995).

15. Jones, G. B. & Trevena, A. J. The influence of coral reefs on atmospheric dimethylsulphide over the Great Barrier Reef, Coral Sea, Gulf of Papua and Solomon and Bismarck Seas. Mar. Freshwater Res. 56, 85–93 (2005).

16. Bourne, D. G., Morrow, K. M. & Webster, N. S. Insights into the Coral Microbiome: Underpinning the Health and Resilience of Reef Ecosystems. Annu. Rev. Microbiol. 70, 317–340 (2016).

17. Pogoreutz, C. et al. The coral holobiont highlights the dependence of cnidarian animal hosts on their associated microbes. Cellular dialogues in the holobiont (2020).

18. Kiene, R. P., Linn, L. J., González, J., Moran, M. A. & Bruton, J. A. Dimethylsulfoniopropionate and Methanethiol Are Important Precursors of Methionine and Protein-Sulfur in Marine Bacterioplankton. Appl Environ Microbiol 65, 4549–4558 (1999).

19. Frade, P. R. et al. Dimethylsulfoniopropionate in corals and its interrelations with bacterial assemblages in coral surface mucus. Environ. Chem. 13, 252–265 (2015).

20. Raina, J.-B., Tapiolas, D., Willis, B. L. & Bourne, D. G. Coral-Associated Bacteria and Their Role in the Biogeochemical Cycling of Sulfur. Applied and Environmental Microbiology 75, 3492–3501 (2009).

21. Tandon, K. et al. Comparative genomics: Dominant coral-bacterium *Endozoicomonas acroporae* metabolizes dimethylsulfoniopropionate (DMSP). ISME J 14, 1290–1303 (2020).

22. Hopkins, F. E., Archer, S. D., Bell, T. G., Suntharalingam, P. & Todd, J. D. The biogeochemistry of marine dimethylsulfide. Nat Rev Earth Environ 4, 361–376 (2023).

23. Zinke, L. Cool gas in warm summers. Nat. Clim. Chang. 9, 434–434 (2019).

24. Vallina, S. M. & Simó, R. Strong Relationship Between DMS and the Solar Radiation Dose over the Global Surface Ocean. Science 315, 506–508 (2007).

25. Reisch, C. R., Moran, M. A. & Whitman, W. B. Bacterial Catabolism of Dimethylsulfoniopropionate (DMSP). Front. Microbio. 2, (2011).

26. Alcolombri, U. et al. Identification of the algal dimethyl sulfide–releasing enzyme: A missing link in the marine sulfur cycle. Science 348, 1466–1469 (2015).

27. Sun, J. et al. The abundant marine bacterium *Pelagibacter* simultaneously catabolizes dimethylsulfoniopropionate to the gases dimethyl sulfide and methanethiol. Nat Microbiol 1, 1–5 (2016).

28. Garren, M. et al. A bacterial pathogen uses dimethylsulfoniopropionate as a cue to target heat-stressed corals. ISME J 8, 999–1007 (2014).

29. Zhang, X. H. et al. Biogenic production of DMSP and its degradation to DMS—their roles in the global sulfur cycle. Science China Life Sciences 62, 1296–1319 (2019).

30. Da Silva, D. M. G., Costa, R. & Keller-Costa, T. A genomic view of the bacterial family Endozoicomonadaceae in marine symbioses. Commun Biol 8, 1418 (2025).

31. Kurahashi, M. & Yokota, A. *Endozoicomonas elysicola* gen. nov., sp. nov., a γproteobacterium isolated from the sea slug Elysia ornata. Systematic and Applied Microbiology 30, 202–206 (2007).

32. Alex, A. & Antunes, A. Comparative Genomics Reveals Metabolic Specificity of *Endozoicomonas* Isolated from a Marine Sponge and the Genomic Repertoire for HostBacteria Symbioses. Microorganisms 7, 635 (2019).

33. Du, Z., Zhang, W., Xia, H., Lü, G. & Chen, G. Isolation and diversity analysis of heterotrophic bacteria associated with sea anemones. Acta Oceanol. Sin. 29, 62–69 (2010).

34. Cano, I. et al. Molecular Characterization of an *Endozoicomonas*-Like Organism Causing Infection in the King Scallop (*Pecten maximus* L.). Applied and Environmental Microbiology 84, e00952–17 (2018).

35. Lim, S. J. et al. Taxonomic and functional heterogeneity of the gill microbiome in a symbiotic coastal mangrove lucinid species. ISME Journal 13, 902–920 (2019).

36. Neave, M. J., Michell, C. T., Apprill, A. & Voolstra, C. R. *Endozoicomonas* genomes reveal functional adaptation and plasticity in bacterial strains symbiotically associated with diverse marine hosts. Sci Rep 7, 40579 (2017).

37. Neave, M. J., Apprill, A., Ferrier-Pagès, C. & Voolstra, C. R. Diversity and function of prevalent symbiotic marine bacteria in the genus *Endozoicomonas*. Appl Microbiol Biotechnol 100, 8315–8324 (2016).

38. Thompson, J. R., Rivera, H. E., Closek, C. J. & Medina, M. Microbes in the coral holobiont: partners through evolution, development, and ecological interactions. Front. Cell. Infect. Microbiol. 4, (2015).

39. Mohamed, A. R., Ochsenkühn, M. A., Kazlak, A. M., Moustafa, A. & Amin, S. A. The coral microbiome: towards an understanding of the molecular mechanisms of coral– microbiota interactions. FEMS Microbiology Reviews 47, fuad005 (2023).

40. Villela, H. et al. Genome analysis of a coral-associated bacterial consortium highlights complementary hydrocarbon degradation ability and other beneficial mechanisms for the host. Sci Rep 13, 12273 (2023).

41. Pratte, Z. A., Stewart, F. J. & Kellogg, C. A. Functional gene composition and metabolic potential of deep-sea coral-associated microbial communities. Coral Reefs 42, 1011–1023 (2023).

42. Hill, R. W. & Dacey, J. W. H. Processing of ingested dimethylsulfoniopropionate by mussels Mytilus edulis and scallops Argopecten irradians. Marine Ecology Progress Series 343, 131–140 (2007).

43. Shu, Y. et al. A bacterial symbiont in the gill of the marine scallop Argopecten irradians irradians metabolizes dimethylsulfoniopropionate. mLife 2, 178–189 (2023).

44. Herry, A., Diouris, M. & Le Pennec, M. Chemoautotrophic symbionts and translocation of fixed carbon from bacteria to host tissues in the littoral bivalve *Loripes lucinalis* (Lucinidae). Marine Biology 101, 305–312 (1989).

45. Brissac, T., Merçot, H. & Gros, O. Lucinidae/sulfur-oxidizing bacteria: ancestral heritage or opportunistic association? Further insights from the Bohol Sea (the Philippines): Hosts/symbionts phylogenetic comparison within Lucinidae. FEMS Microbiology Ecology 75, 63–76 (2011).

46. Childress, J. J. & Girguis, P. R. The metabolic demands of endosymbiotic chemoautotrophic metabolism on host physiological capacities. Journal of Experimental Biology 214, 312–325 (2011).

47. Dubilier, N., Bergin, C. & Lott, C. Symbiotic diversity in marine animals: the art of harnessing chemosynthesis. Nature Reviews Microbiology 6, 725–740 (2008).

48. Petersen, J. M. et al. Chemosynthetic symbionts of marine invertebrate animals are capable of nitrogen fixation. Nat Microbiol 2, 16195 (2016).

49. Taylor, J. D. & Glover, E. Biology, Evolution and Generic Review of the Chemosymbiotic Bivalve Family Lucinidae. (Ray Society, London, 2021).

50. Williams, B. T. et al. Bacteria are important dimethylsulfoniopropionate producers in coastal sediments. Nature Microbiology 4, 1815–1825 (2019).

51. Lim, S. J. et al. Extensive Thioautotrophic Gill Endosymbiont Diversity within a Single *Ctena orbiculata* (Bivalvia: Lucinidae) Population and Implications for Defining HostSymbiont Specificity and Species Recognition. mSystems 4, 1–19 (2019).

52. Curson, A. R. J. et al. Dimethylsulfoniopropionate biosynthesis in marine bacteria and identification of the key gene in this process. Nature Microbiology 2, (2017).

53. Curson, A. R. J., Fowler, E. K., Dickens, S., Johnston, A. W. B. & Todd, J. D. Multiple DMSP lyases in the γ-proteobacterium *Oceanimonas doudoroffii*. Biogeochemistry 110, 109–119 (2012).

54. Sudo, M. L. Host-microbe-environment interactions in sulphur-oxidising symbioses. (University of Vienna, 2023).

55. Sudo, M. et al. *SoxY* gene family expansion underpins adaptation to diverse hosts and environments in symbiotic sulfide oxidizers. mSystems 9, e01135–23 (2024).

56. Pjevac, P. et al. An Economical and Flexible Dual Barcoding, Two-Step PCR Approach for Highly Multiplexed Amplicon Sequencing. Frontiers in Microbiology 12, (2021).

57. Herlemann, D. P. et al. Transitions in bacterial communities along the 2000 km salinity gradient of the Baltic Sea. ISME J 5, 1571–1579 (2011).

58. Pruesse, E. et al. SILVA: a comprehensive online resource for quality checked and aligned ribosomal RNA sequence data compatible with ARB. Nucleic Acids Research 35, 7188–7196 (2007).

59. Pruesse, E., Peplies, J. & Glöckner, F. O. SINA: Accurate high-throughput multiple sequence alignment of ribosomal RNA genes. Bioinformatics 28, 1823–1829 (2012).

60. Kolmogorov, M., Yuan, J., Lin, Y. & Pevzner, P. A. Assembly of long, error-prone reads using repeat graphs. Nat Biotechnol 37, 540–546 (2019).

61. Morel-Letelier, I. et al. Adaptations to nitrogen availability drive ecological divergence of chemosynthetic symbionts. PLoS Genet 20, e1011295 (2024).

62. Osvatic, J. T., et al. Global biogeography of chemosynthetic symbionts reveals both localized and globally distributed symbiont groups. Proc. Natl. Acad. Sci. U.S.A. 118, e2104378118 (2021).

63. Parks, D. H., Imelfort, M., Skennerton, C. T., Hugenholtz, P. & Tyson, G. W. CheckM: assessing the quality of microbial genomes recovered from isolates, single cells, and metagenomes. Genome Res. 25, 1043–1055 (2015).

64. Chaumeil, P.-A., Mussig, A. J., Hugenholtz, P. & Parks, D. H. GTDB-Tk: a toolkit to classify genomes with the Genome Taxonomy Database. Bioinformatics 36, 1925–1927 (2020).

65. Minh, B. Q. et al. IQ-TREE 2: New Models and Efficient Methods for Phylogenetic Inference in the Genomic Era. Molecular Biology and Evolution 37, 1530–1534 (2020).

66. Hoang, D. T., Chernomor, O., von Haeseler, A., Minh, B. Q. & Vinh, L. S. UFBoot2: Improving the Ultrafast Bootstrap Approximation. Molecular Biology and Evolution 35, 518–522 (2018).

67. Bianchini, G. & Sánchez-Baracaldo, P. TREEVIEWER : Flexible, modular software to visualise and manipulate phylogenetic trees. Ecology and Evolution 14, e10873 (2024).

68. Jain, C., Rodriguez-R, L. M., Phillippy, A. M., Konstantinidis, K. T. & Aluru, S. High throughput ANI analysis of 90K prokaryotic genomes reveals clear species boundaries. Nat Commun 9, 5114 (2018).

69. Tanabe, T. S. & Dahl, C. HMSS2 : An advanced tool for the analysis of sulphur metabolism, including organosulphur compound transformation, in genome and metagenome assemblies. Molecular Ecology Resources 1755–0998.13848 (2023).

70. Camacho, C. et al. BLAST+: architecture and applications. BMC Bioinformatics 10, 421 (2009).

71. Gertz, E. M., Yu, Y.-K., Agarwala, R., Schäffer, A. A. & Altschul, S. F. Compositionbased statistics and translated nucleotide searches: Improving the TBLASTN module of BLAST. BMC Biology 4, 41 (2006).

72. Aziz, R. K. et al. The RAST Server: Rapid Annotations using Subsystems Technology. BMC Genomics 9, 75 (2008).

73. Brettin, T. et al. RASTtk: A modular and extensible implementation of the RAST algorithm for building custom annotation pipelines and annotating batches of genomes. Sci Rep 5, 8365 (2015).

74. Overbeek, R. et al. The SEED and the Rapid Annotation of microbial genomes using Subsystems Technology (RAST). Nucleic Acids Research 42, D206–D214 (2014).

75. Mahram, A. & Herbordt, M. C. NCBI BLASTP on high-performance reconfigurable computing systems. ACM Transactions on Reconfigurable Technology and Systems (TRETS) 7, 1–20 (2015).

76. Jones, P. et al. InterProScan 5: Genome-scale protein function classification. Bioinformatics 30, 1236–1240 (2014).

77. Finn, R. D., Clements, J. & Eddy, S. R. HMMER web server: interactive sequence similarity searching. Nucleic Acids Research 39, W29–W37 (2011).

78. Wang, J. et al. Alternative dimethylsulfoniopropionate biosynthesis enzymes in diverse and abundant microorganisms. Nat Microbiol 9, 1979–1992 (2024).

79. Sievers, F. et al. Fast, scalable generation of high-quality protein multiple sequence alignments using Clustal Omega. Molecular Systems Biology 7, 539 (2011).

80. Hill, R. W. & Dacey, J. W. H. Exceptional accumulation and retention of dimethylsulfoniopropionate by molluscs. Environ. Chem. 13, 231–238 (2015).

81. Van Der Geest, M. et al. Nutritional and reproductive strategies in a chemosymbiotic bivalve living in a tropical intertidal seagrass bed. Marine Ecology Progress Series 501, 113–126 (2014).

82. Alcaraz, C. M. et al. Sulfur-oxidizing symbionts colonize the digestive tract of their lucinid hosts. The ISME Journal 18, wrae200 (2024).

83. Smit, A. J., Robertson-Andersson, D. V., Peall, S. & Bolton, J. J. Dimethylsulfoniopropionate (DMSP) accumulation in abalone Haliotis midae (Mollusca: Prosobranchia) after consumption of various diets, and consequences for aquaculture. Aquaculture 269, 377–389 (2007).

84. Kocsis, M. G. et al. Dimethylsulfoniopropionate biosynthesis in Spartina alterniflora 1: evidence that S-methylmethionine and dimethylsulfoniopropylamine are intermediates. Plant Physiology 117, 273–281 (1998).

85. Ball, A. D., Purdy, K. J., Glover, E. A. & Taylor, J. D. Ctenidial structure and three bacterial symbiont morphotypes in Anodontia (Euanodontia) ovum (Reeve, 1850) from the Great Barrier Reef, Australia (Bivalvia: Lucinidae). Journal of Molluscan Studies 75, 175–185 (2009).

86. Beinart, R. A., Nyholm, S. V., Dubilier, N. & Girguis, P. R. Intracellular O ceanospirillales inhabit the gills of the hydrothermal vent snail *Alviniconcha* with chemosynthetic, γ-P roteobacterial symbionts. Environ Microbiol Rep 6, 656–664 (2014).

87. Chiou, Y.-J. et al. Similar but different: Characterization of dddD gene–mediated DMSP metabolism among coral-associated Endozoicomonas. Science Advances 9, eadk1910 (2023).

88. Pogoreutz, C. et al. Coral holobiont cues prime *Endozoicomonas* for a symbiotic lifestyle. ISME J 16, 1883–1895 (2022).

89. Sun, L., Curson, A. R. J., Todd, J. D. & Johnston, A. W. B. Diversity of DMSP transport in marine bacteria, revealed by genetic analyses. Biogeochemistry 110, 121–130 (2012).

90. Todd, J. D. et al. Structural and Regulatory Genes Required to Make the Gas Dimethyl Sulfide in Bacteria. Science 315, 666–669 (2007).

91. Todd, J. D. et al. Molecular dissection of bacterial acrylate catabolism - unexpected links with dimethylsulfoniopropionate catabolism and dimethyl sulfide production. Environmental Microbiology 12, 327–343 (2010).

92. Carrión, O. et al. DMSOP-cleaving enzymes are diverse and widely distributed in marine microorganisms. Nat Microbiol 8, 2326–2337 (2023).

93. Liu, J. et al. Novel Insights Into Bacterial Dimethylsulfoniopropionate Catabolism in the East China Sea. Front Microbiol 9, 3206 (2018).

94. Eyice, Ö. et al. Bacterial SBP56 identified as a Cu-dependent methanethiol oxidase widely distributed in the biosphere. ISME Journal 12, 145–160 (2018).

95. Pol, A., Op Den Camp, H. J. M., Mees, S. G. M., Kersten, M. A. S. H. & Van Der Drift, C. Isolation of a dimethylsulfide-utilizing Hyphomicrobium species and its application in biofiltration of polluted air. Biodegradation 5, 105–112 (1994).

96. Carrión, O. et al. Methanethiol-dependent dimethylsulfide production in soil environments. The ISME Journal 11, 2379–2390 (2017).

97. Stipanuk, M. H. Metabolism of Sulfur-Containing Amino Acids: How the Body Copes with Excess Methionine, Cysteine, and Sulfide. The Journal of Nutrition 150, 2494S2505S (2020).

98. Tuite, N. L., Fraser, K. R. & O’Byrne, C. P. Homocysteine Toxicity in *Escherichia coli* Is Caused by a Perturbation of Branched-Chain Amino Acid Biosynthesis. J Bacteriol 187, 4362–4371 (2005).

99. Manzano-Marín, A. et al. Co-obligate symbioses have repeatedly evolved across aphids, but partner identity and nutritional contributions vary across lineages. Peer Community Journal 3, e46 (2023).

100. McCutcheon, J. P., McDonald, B. R. & Moran, N. A. Convergent evolution of metabolic roles in bacterial co-symbionts of insects. Proc. Natl. Acad. Sci. U.S.A. 106, 15394–15399 (2009).

