## Supplementary for "Organosulfur cycling in the multi-partner symbiosis between lucinid bivalves, sulphur-oxidizing symbionts, and *Endozoicomonas*"

### Supplementary Information

Additional supplementary files:

Methods: Supplementary table SM1- List of genomes used in the study/phylogeny

Supplementary table SM2 – List of genes used for blast search

Results: Supplementary table SR2 - Gene presence/absence table

Supplementary table SR3 – CheckM table

Supplementary table SR4 – ANI scores

#### SI Methods

##### *Sampling of* Loripes orbiculatus *and* Loripinus fragilis *for molecular work*

The DNA for 16S rRNA gene amplicon sequencing was extracted from *Loripes orbiculatus* and *Loripinus fragilis* gills (9 samples each) obtained during environmental sampling and laboratory incubation. Clams were collected within a *Cymodocea nodosa* seagrass meadow using sediment extraction with metal corer with big diameter of 30 cm and sieving. Species identification was based on morphological characteristics. A subset of the collected clams was immediately processed - these represent 'natural' conditions at the time of sampling. The remaining clams (three of each species) were placed in a glass beaker with site seawater overnight before the start of the incubations. The incubation vials (50 ml, 25×150 mm, culture tubes with screw cap, KIMAX®, KIMBLE®) contained a sulphide agar plug (2% agar and 5 mM of sodium sulphide) overlaid with 5 cm of acid-cleaned glass beads (0.75-1 mm, Sigma-Aldrich), and filled with seawater from the sampling site. The vials were incubated on ice overnight to establish an equilibrium and create a sulphide and oxygen gradient mimicking a marine sediment (Figure 1b). After 15 h, the incubation vials were transferred to room temperature (RT). The incubations were done at RT for 24h. Immediately after sieving or at the end of the incubation period, clams' gills were dissected, fixed in RNAlater (Thermo Fisher Scientific) and stored overnight at 4°C and subsequently at − 80 °C. The incubations were not expected to change the composition of microbial community, as reflected in the data. For the assembly of de novo transcriptomes of the *L. orbiculatus* and *L. fragilis* we used only RNA from freshly collected clams.

##### *DNA and RNA extractions and processing*

The extraction of nucleic acids from gills was performed according to the TRIzol™ (Thermo Fischer Scientific) extraction protocol, which was adjusted for clam tissue with the following minor modifications: the protocol recommended for small quantities of the tissue was used, bromochloropropane (BCP) was used for the initial phase separation, and phase separation for DNA extraction was carried performed using the back extraction buffer (4 M guanidine thiocyanate, 50 mM sodium citrate and 1 M Tris) (see protocol below). The homogenisation was performed by dicing the gill and manually grinding it using tissue grinders and glass beads. The extracted RNA was quantified with the Qubit 4 Fluorometer (Thermo Fischer Scientific) using a Qubit RNA BR Assay Kit (Thermo Fischer Scientific) according to the manufacturer’s protocol. The DNA contamination was determined by PCR 16 rRNA gene with 616V (5'- AGA GTT TGA TYM TGG CTC AG-3') and 1492R (5'-GGT TAC CTT GTT ACG ACT T-3') primers and gel electrophoresis as described in “Isolation and cultivation of *Ca*. E. endolucinida” section below. The samples, which were contaminated with DNA, underwent treatment with the Turbo DNA-free Kit (Thermo Fisher Scientific), according to the manufacturer's instructions. The quality of extracted RNA was assessed using a NanoDrop (Nanodrop Technologies, Thermo Scientific, Waltham, MA, United States) and High Sensitivity RNA ScreenTape® on TapeStation 4200 (Agilent). The extracted DNA was quantified with the Qubit 4 Fluorometer (Thermo Fischer Scientific) using a Qubit DNA BR Assay Kit (Thermo Fischer Scientific) according to the manufacturer’s protocol.

**Trizol DNA/RNA extraction protocol (bead beating) adapted from F_071215 Invitrogen Trizol detailed protocol**

The TRIzol™ (Thermo Fischer Scientific) extraction protocol was used with minor modifications for clam tissue. The recommended protocol for small quantities of the tissue was followed, and bromochloropropane (BCP) was used for the initial phase separation. The back extraction buffer (4 M guanidine thiocyanate, 50 mM sodium citrate, and 1 M Tris) was used for phase separation during DNA extraction.

Homogenization using bead beater

1. Add gill sample to bead beater lysis tube (lysis matrix D)
2. Add 990µl Trizol
3. Add 12.5µl glycogen (20mg/mL)
4. Put tubes into bead beater for 10 s, 4.0 m/s
5. Incubate at RT for 5 min

Phase separation #1

6. 1. Spin at max speed for 10 mins, 4°C
7. 2. Transfer supernatant into the new Eppendorf tube
8. 3. Add 160µl 1-bromo-3-chloropropane (BCP)
9. 4. Invert or shake for 15 s and incubate at RT for 15 min
10. 5. Spin at max speed for 15 min, 4°C. Three layers should be visible
11. 6. Transfer colourless aqueous phase (RNA) into a low bind Eppendorf tube
12. Re-spin 5 min at max speed if layers slip

Phase separation #2

13. Remove remaining aqueous phase
14. Add 0.5 initial-Trizol volume (500µl to 1ml) of back extraction buffer (4M guanidine thiocyanate, 50mM sodium citrate, 1M Tris pH 8.0)
15. Shake tubes vigorously by hand for 15 s
16. Incubate 10 min at RT
17. Phase separation: centrifuge samples 12 000 g (10 620 rpm*), 15 min, 4°C
18. Transfer the aqueous phase containing DNA to a new Eppendorf tube

DNA precipitation

19. Add 12.5µl glycogen (20mg/mL)
20. Add 0.4 ml of isopropanol per 1 ml of Trizol used initially (400µl per 1ml Trizol)
21. Mix by inversion and incubate at -80°C for 1 hour – overnight

DNA wash and elution

10. Spin 12000 x g (10 620 rpm*) for 5 min at 4-25°C and remove the supernatant
11. Add 1 ml of 0.1 M sodium citrate in 10% ethanol (pH 8.5) to the DNA pellet, mix by inversion, incubate for 30 min at room temperature
12. Centrifuge samples 4 000 g for 5 min 4°C, discard supernatant
13. Wash by adding 1 ml 75% EtOH, incubate for 15 min at room temperature
14. Centrifuge samples 4 000 g for 5 min 4°C to pellet the DNA
15. Discard the supernatant
16. Remove ethanol, re-spin a few seconds and remove remaining ethanol
17. Air dry the DNA pellet for 5-10 min, but do not allow the pellet to dry out
18. Add 70 µl of nuclease-free water (pre-heat to 65°C), TE per 10 mg of tissue (final conc. should be 0.2-0.3 µg/µl) by slow and gentle pipetting.

rpm values for Eppendorf microcentrifuge 5430R (FA-45-30-11, Eppendorf SE, Germany)

*RNAseq: Library construction and sequencing, sequence pre-processing and filtering*

In all samples, rRNA was depleted using Illumina's Ribo Zero Plus rRNA depletion kit following the manufacturer's protocol. Sequencing libraries were prepared from the rRNA-depleted RNA with the NEBNext® Ultra™ II Directional RNA Library Prep Kit for Illumina and sequenced on the NovaSeq6000 (SP, 2x 100 bp mode) at the Biomedical Sequencing Facility (BSF4, Vienna. Bam files were converted to fastq format using samtools v. 1.18. [1]. Predicted errors in the raw reads were corrected using Rcorrector v 1.0.5 under default settings, and the uncorrectable reads were removed [2]. The script FilterUncorrectabledPEfastq.py, a part of TranscriptomeAssemblyTools (https://github.com/harvardinformatics/TranscriptomeAssemblyTools), has been used to remove broken reads. Further, the reads were trimmed from adapters, and low-quality reads were removed using TrimGalore v. 0.6.10, a wrapper script around Cutadapt [3]. To eliminate rRNA sequences from the libraries, we used SortMeRNA v. 4.3.6, which searched against the default rRNA database of the package that includes 5S, 5.8S, 16S, 23S, 18S, and 28S rRNA sequences [4]. For small transformations such as extension conversion or split into paired files, bbmap-39.01 (sourceforge.net/projects/bbmap/) and Seqkit-2.5.1 [5] were used. The quality of the processed reads has been analysed using FastQC v0.12.1 [6]. Based on the FastQC results, overrepresented sequences have been removed using RemoveFastqcOverrepSequenceReads.py script from TranscriptomeAssemblyTools. The final quality check has been performed using FastQC v0.12.1 and multiqc-1.15 [7].

*De novo transcriptome assembly of lucinid hosts and quality assessment*

To assemble the host transcriptome, we first cleaned the total reads of any symbiont sequences by mapping them to best-quality genome of each of the (dominant) symbiont species *Ca.* Sedimenticola endoloripinus or *Ca.* Thiodiazotropha luna using bwa-mem function of BWA software package [8]. The transcriptome reads which didn't map to the symbiont genome were extracted with samtools v. 1.18. [1].

The resulting bam files were converted to fastq files using the bamToFastq option within bedtools v2.30.0 [9]. For each of the hosts, all reads have been concatenated, and missing paired reads repaired using bbmap v. 39.01 [10]. Transcripts have been assembled using Trinity v. 2.15.1 and the standard protocol [11,12]. The duplicates were removed using the cd-hit-est function of cdhit v. 4.8.1 with a sequence identity threshold of 100% [13]. Initial quality control was performed using BUSCO v. 5.5.0 [14]. We used transdecoder v. 5.7.0 to identify candidate coding regions from the assembled transcripts with a minimum length of 100. We ran TransDecoder.Predict with the "--single_best_only" option so that only the single best candidate region per transcript was considered for further analysis.

*DSYB candidate search and structural prediction*

To investigate the genomic potential of DMSP production by animal host, we searched the de novo assembled transcriptomes of *Loripes orbiculatus* and *Loripinus fragilis* for DSYB homologs based on hmm profiles and amino acid sequenced of several DMSP synthesizing genes (main manuscript). Due to the low amino acid identity, we further investigated the putative DSYB sequences from lucinid hosts by InterProScan Search [15] and AlphaFold prediction via ColabFold [16] and visualisation within ChimeraX[17].

*Isolation and cultivation of* Ca. *Endozoicomonas endolucinida*

The 16S rRNA genes were amplified using the primers 616V (5'- AGA GTT TGA TYM TGG CTC AG-3') and 1492R (5'-GGT TAC CTT GTT ACG ACT T-3'). The colony PCR was preceded by a pre-lysis step where a single colony was picked from the plate and dissolved in 20 µl of MQ water, frozen at -20°C for 30 min and incubated at 95°C for 10 min. Lysed cells were centrifuged at 15.000 g, and 10 µl of the supernatant was added to the PCR reaction mix. The PCR reaction mix (final concentrations: Green 1X Dream Taq Buffer, dNTPS 0.2 mM, BSA 0.2 mg/ml, Taq polymerase 0.05 U/µl, primers 1 µM) was prepared in a final volume of 50 µl per reaction. The amplification cycles were as follows: initial denaturation at 95 °C for 3 min, followed by 30 cycles at 95 °C for 30 s, 56 °C for 30 s, 72 °C for 1.5 min, and a final elongation at 72 °C for 10 min. PCR products were visualised on 2% agarose gel electrophoresis and subsequently purified using a QIAquick PCR Purification Kit (Qiagen) according to the manufacturer's instructions. Concentrations were measured by Qubit 4 Fluorometer (Thermo Fischer Scientific) using a Qubit DNA BR Assay Kit (Thermo Fischer Scientific) according to the manufacturer's protocol. Samples were sent for Sanger sequencing at Microsynth AG (Vienna, Austria).

*DMSP uptake by Ca.* Endozoicomonas endolucinida

To measure DMSP levels in both the cell pellet and dissolved supernatant fractions. Samples culture were centrifuged to separate the cell pellet and the culture supernatant. The pellet fraction was resuspended in 0.2 mL PBS buffer and treated with 100 µL NaOH (10 M) to chemically cleave the DMSP and yield DMS. The cell-free supernatant (200 µL) was similarly treated for DMS analysis. The results are reported as DMSPp and DMSPd.

*Phylogenetic classification of Endozoicomonas sp.*

16S rRNA gene sequences were annotated in the isolated genome using BARRNAP v0.9. These sequences were combined with all 16S rRNA genes obtained from the culture of isolated bacteria from Sanger sequencing using the 1492r primer and with ASVs assigned as *Endozoicomonas* during the amplicon data analysis. 16S rRNA gene sequences were aligned and taxonomically classified using the SILVA: SINA web server under default parameters and the "Search and classify" workflow [18,19]. Furthermore, the 16S rRNA genes of *Endozoicomonas acroporae* (SILVA ID: PJPV01000151.193.1576), used in this study as a representative of DMSP degrading *Endozoicomonas*, as well as of *Marinobacter hydrocarbonoclasticus* (SILVA ID: AB021372.1.1512) were used as an outgroup accordingly to [20]. We also included previously reported Endozoicomonas OTUs with NCBI IDs: KY687503.2 and KY687505.3 occurring in lucinid clam *Ctena orbiculata* [21]. The alignment was enriched with ten neighbours per query sequence. The alignment was used to generate a phylogenetic tree calculated with the "RAxML" method.

*Live incubations of* Loripes orbiculatus

Clams were incubated in airtight 50 ml serum vials, open to air in oxic, sealed in anoxic incubations. We used 50 ml of 0.2 µm filtered seawater from the sampling site in Piran for each incubation vial and added antibiotics (Gentamicin, Chloramphenicol, and Ampicillin at 100 µg/ml dissolved in water and A22 at 20 µg/ml dissolved in DMSO) in respective treatments. We added 50 µl of DMSO to the control treatment to ensure the same amount of potential substrates for DMS or DMSP production. We measured oxygen concentrations during incubations using a fibre-optic oxygen sensor (OXR430, PyroScience, Germany) at the bottom of the flask after 6, 23, 30, 33, and 48 hours from the beginning of the incubations. The oxygen sensor was calibrated using 0% $O_2$ Calibration Capsules (OXCAL, PyroScience, Germany). After the experiment, we carefully dissected the clams and prepared them for DMSP measurements via the alkaline lysis and GC method described above.

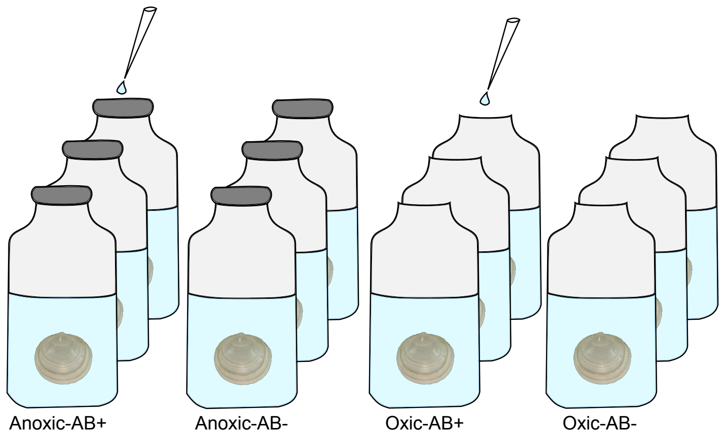


Figure SM1: Schematic representation of experimental setup, which includes four treatments: anoxic control and with the addition of antibiotics and oxic control or with the addition of antibiotics. Each of the treatments was done in triplicates.

### SI Results and discussion

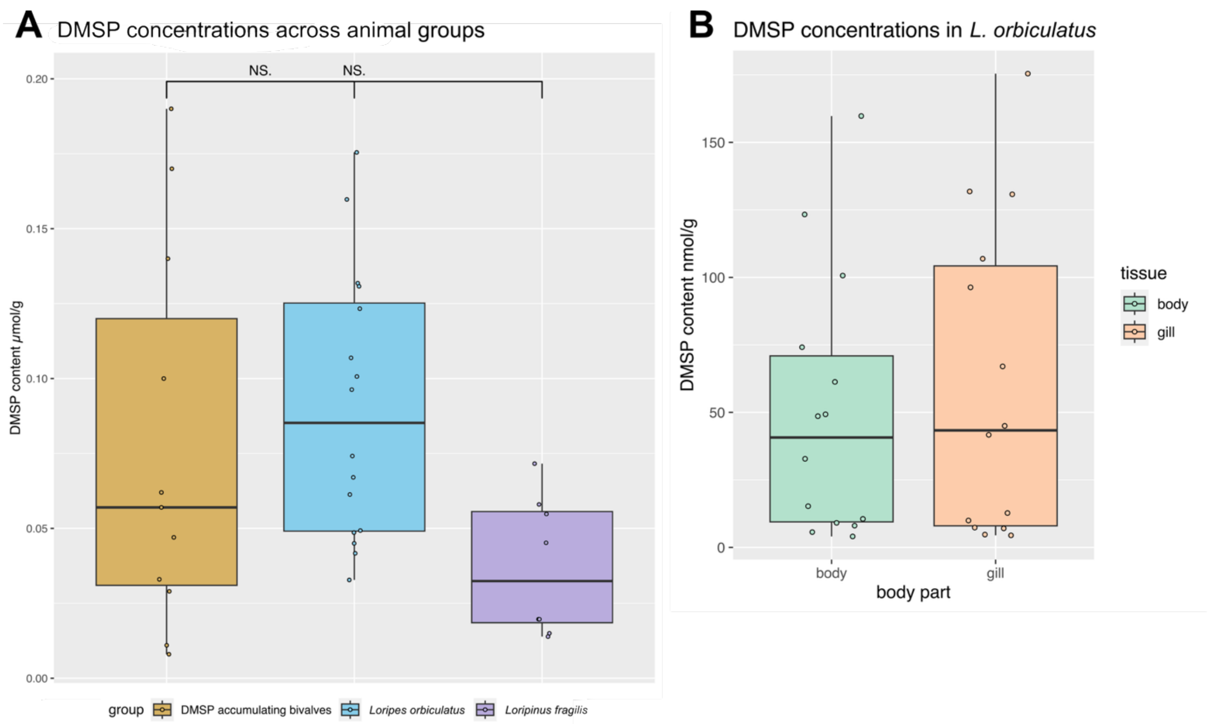


Figure SR1: (A) DMSP content in *Loripes orbiculatus* and *Loripinus fragilis* compared to non-symbiotic DMSP accumulating bivalve from Hill & Dacey 2007 [22]. None of the groups showed significant differences between each other. (B) DMSP content in different tissue types of *Loripes orbiculatus* across all samples. The DMSP values measured in the symbiont harbouring gills are not significantly different from the symbiont-free body suggesting symbiont independent DMSP production or accumulation.

DMSP content in lucinid clams is not significantly different from DMSP accumulating bivalves *Mytilus edulis* and *Argopecten irradians,* the second of which is known to harbour DMSP-degrading symbiotic bacteria. DMSP values in all measured lucinids show high standard deviation, which is likewise found in the DMSP-accumulating molluscs shown in the Figure SR1 as well as in *Geukensia demissa* and *Mytilus edulis* [23].

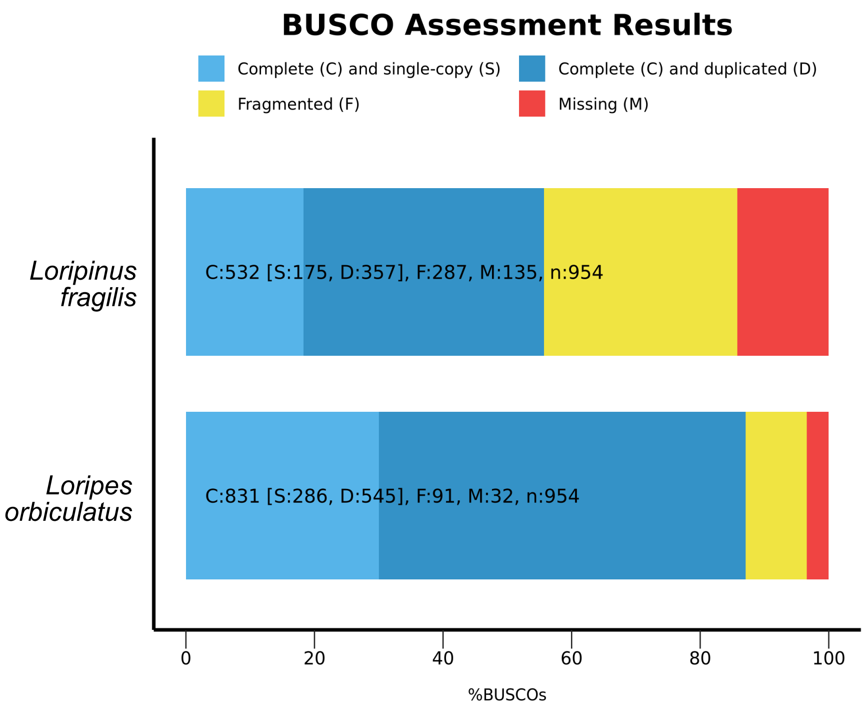


Figure SR2: BUSCO assessment results for de novo assembled transcriptomes of *Loripes orbiculatus* and *Loripinus fragilis*. Despite a higher completeness of *L. orbiculatus*, both transcriptomes lack a substantial number of genes which may have influenced the DSYB homolog search.

| | |
|---|---|
| **Read N50 (bp)** | 5862328 |
| **# contigs** | 4 |
| **Assembly size (bp)** | 6422439 |
| **N50 (bp)** | 5862328 |
| **GC (%)** | 50.1% |
| **Largest contig (bp)** | 5862328 |
| **Completeness** | 98.28 % |
| **Contamination** | 1,89 |
| **Strain heterogeneity** | 12,5 |
| **Kingdom_GTDB** | d__Bacteria |
| **Phylum_GTDB** | p__Pseudomonadota |
| **Class_GTDB** | c__Gammaproteobacteria |
| **Order_GTDB** | o__Pseudomonadales |
| **Family_GTDB** | f__Endozoicomonadaceae |
| **Genus_GTDB** | g__Endozoicomonas |
| **Species_GTDB** | s__ |

Table SR1: Statistics of *Endozoicomonas* sp. isolate assembly.

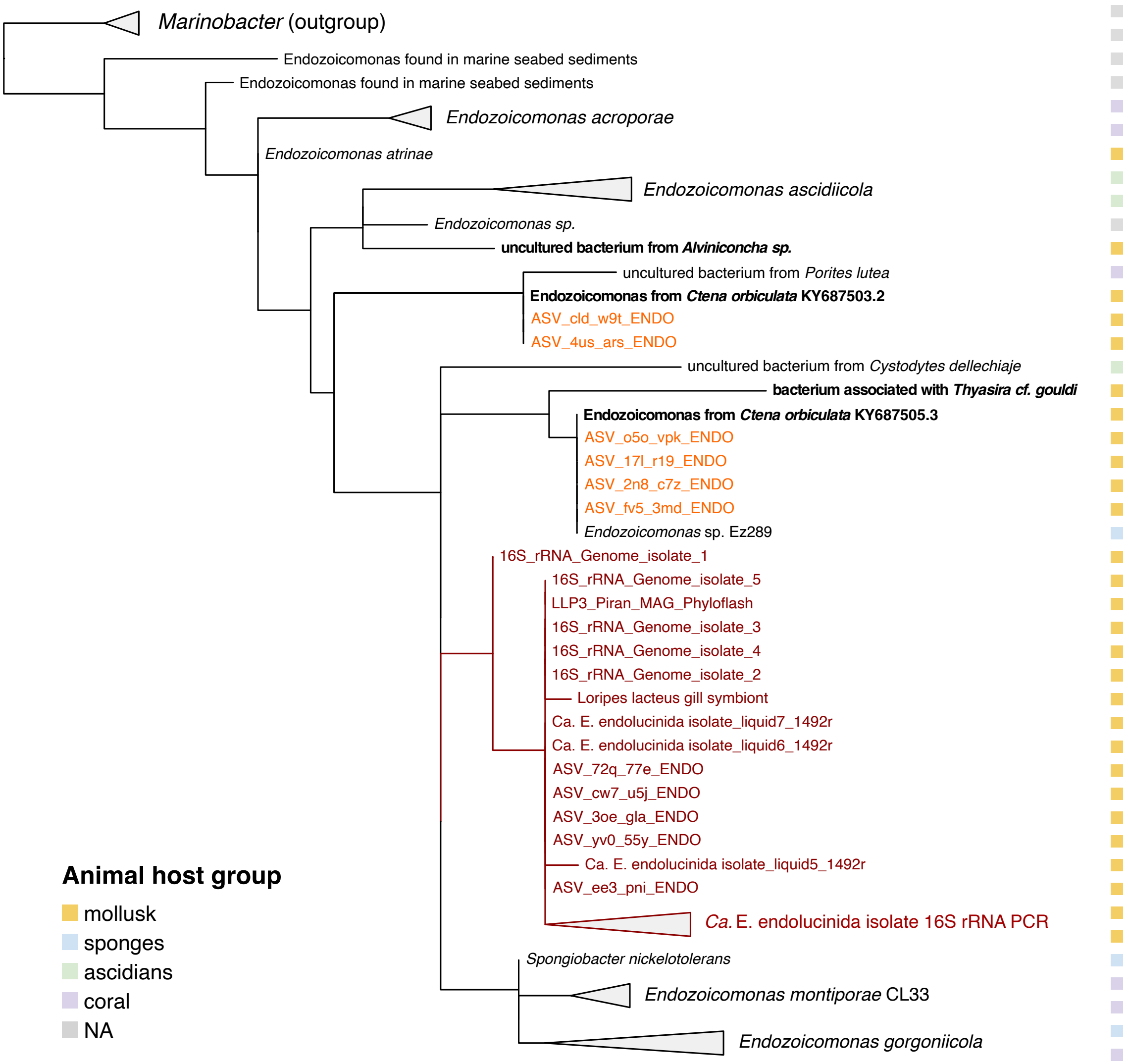


Figure SR3: Phylogenetic relationships of 16S rRNA gene using randomised accelerated maximum likelihood according to the SILVA-SINA workflow. Red colour indicates sequences of *Ca*. E. endolucinida, orange indicates other *Endozoicomonas* ASVs found in Lucinids investigates in this study, *Endozoicomonas* found in other sulphur-oxidizing symbioses are marked as bold. Colours on the side of the tree indicate the animal group in which *Endozoicomonas* was found, according to the legend. The tree was rooted based on *Marinobacter* as an outgroup.

Based on the phylogenetic analysis of the 16S rRNA gene, the *Endozoicomonas* isolates in this study, along with the majority of *Endozoicomonas* ASVs identified in lucinid gills, show close relatedness to *Endozoicomonas elysicola,* a bacterium originally isolated from the gastrointestinal tract of the sea slug *Elysia ornata* [24]. Additionally, all five 16S rRNA gene sequences obtained from the genome of the *Endozoicomonas* sp. isolated in our study cluster with sequences obtained through colony PCR, sanger sequencing, and the five recovered lucinid gill ASVs. This clustering suggests that these sequences collectively represent the same bacterial species. Notably, a sequence from the SILVA database labelled "*Loripes lacteus* gill symbiont" is also present in this clade, representing an *Endozoicomonas* 16S rRNA gene identified in *Loripes orbiculatus* sampled previously in Croatia [25].

Further ASVs identified in lucinid gills in this study cluster with various undescribed *Endozoicomonas* obtained from corals, sponges, and other molluscs. Notably, none of the *Endozoicomonas* sp. sequences from the lucinid clams group with *Endozoicomonas acroporae*, a species confirmed to degrade host-produced DMSP in corals.  However, two ASVs (ASV_cld_w9t and ASV_4us_ars) are found in a clade with *Endozoicomonas* from the coral *Porites lutea*. Additionally, two sequences from *Endozoicomonas* associated with molluscs hosting sulphur-oxidising symbionts, *Thyasira* cf. *gouldi* and *Alvinoconca sp*., were identified during a SILVA search. Notably, the *Endozoicomonas* from *Alvinoconca sp.* forms a sister lineage to *E. acroporae*, presenting an intriguing aspect of the evolutionary relationships within this bacterial group.

##### *DMSP uptake and production by Ca*. Endozoicomonas endolucinida

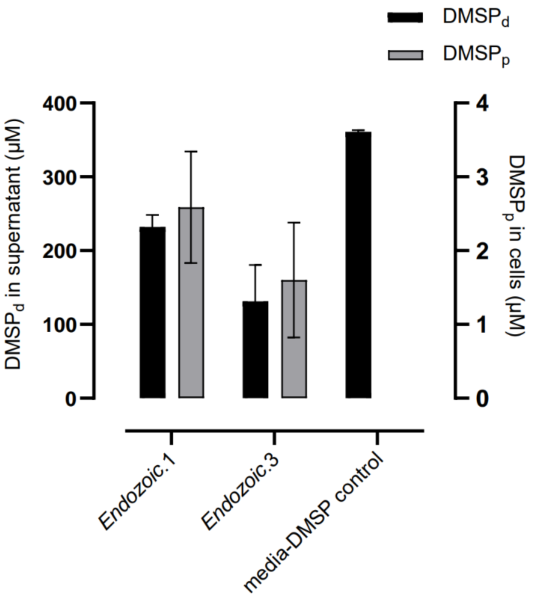


Figure SR4: DMSP uptake by *Ca*. Endozoicomonas endolucinida isolates. Both isolates 1 and 3 incubated in DMSP showed measurable particulate (DMSPp, grey) levels and decreased dissolved (DMSPd, black) levels in the media after incubation. There was no DMSPp detectable in the bacteria-free media.

We were able to measure uptake of DMSP and conversion into DMSPp by our Endozoicomonas isolate. We did not find any genomic DMSP production potential (Supplementary table SR2) or DMSP production activity (data not shown).

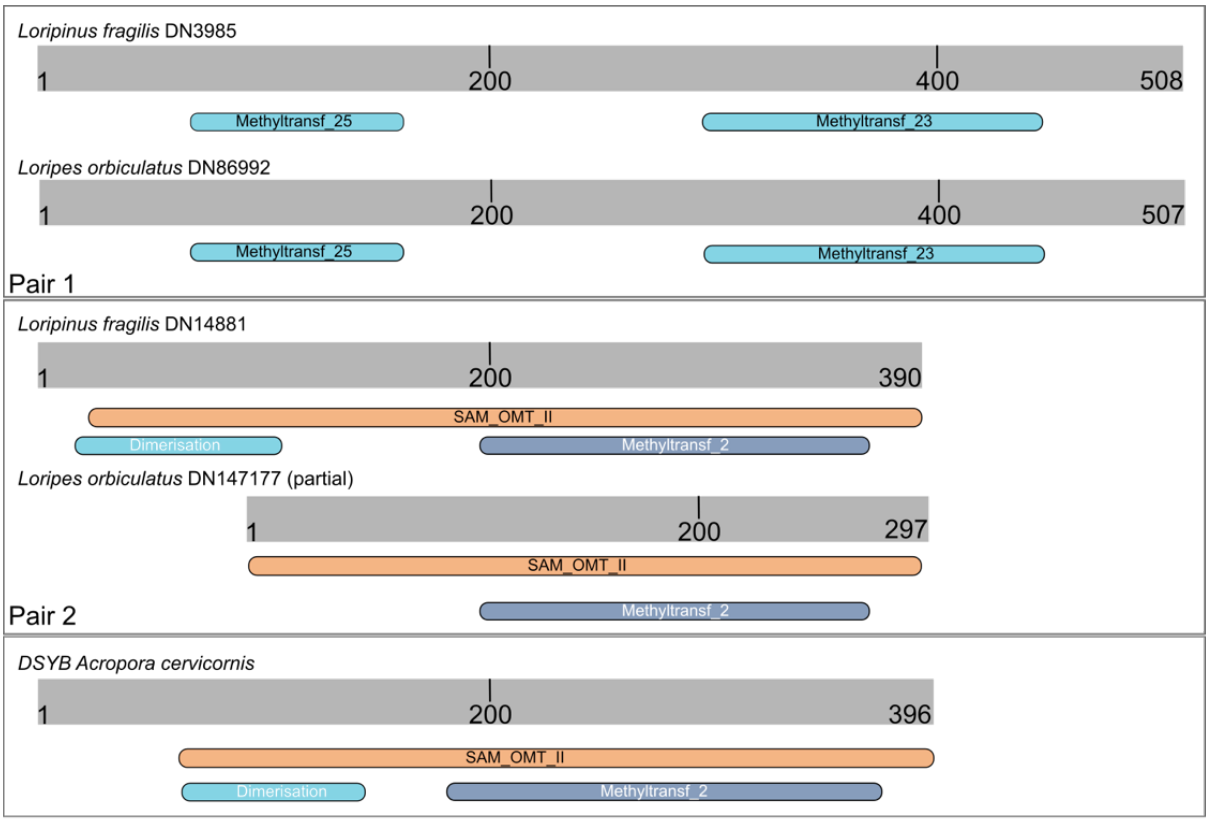


Figure SR5: InterProScan protein domains for candidate DSYBs in *Loripes orbiculatus* and *Loripinus fragilis* clams and DSYB described in coral *Acropora cervicornis*. The length of sequences from the second pair of putative lucinid DSYBs of 390 aa resembles more the typical DSYB aa-length than the first pair of putative lucinid DSYBs with length of 509 aa. Pair 1 DSYB candidates has two domains: Methyltransf_23 (PF13489) and Methyltransf_25 (PF13649 or IPR041698). Pair 2 DSYB candidates from lucinid clams have the same protein domains as DSYB candidate in *Acropora cervicornis*, i.e. the O- methyltransferase domain, also known as Methyltransf_2 (PF00891 or IPR001077) also classified as S-adenosylmethionine-dependent methyltransferases domain (SAM or AdoMet-MTase), class I (cd02440) as well as O-methyltransferase dimerisation domain (PF08100 or IPR012967) assigning this peptide as SAM-dependent O-methyltransferase class II-type profile (PS51683 or IPR016461).

The candidate genes (DN3985 and DN86992) showed the highest aa identity and lowest e-value with DSYE homologs (27.5% and e-value of 7.54E-19; 27.5% and e-value of 1.81E-18 respectively), but clustered on the phylogenetic three with phosphatidylethanolamine N-methyltransferases (pmt), including sequences from the nematode *Caenorhabditis elegans* and *Arabidopsis thaliana* (Figure 3A). These candidates lacked the O-methyltransferase domain (PF00891), typically reported in DSYB (Supplementary figure SR5). In other bivalves, phosphatidylethanolamine N-methyltransferases catalyse phosphatidylcholine synthesis for the biosynthesis of betaine, an organic osmolyte in marine animals [26–28].

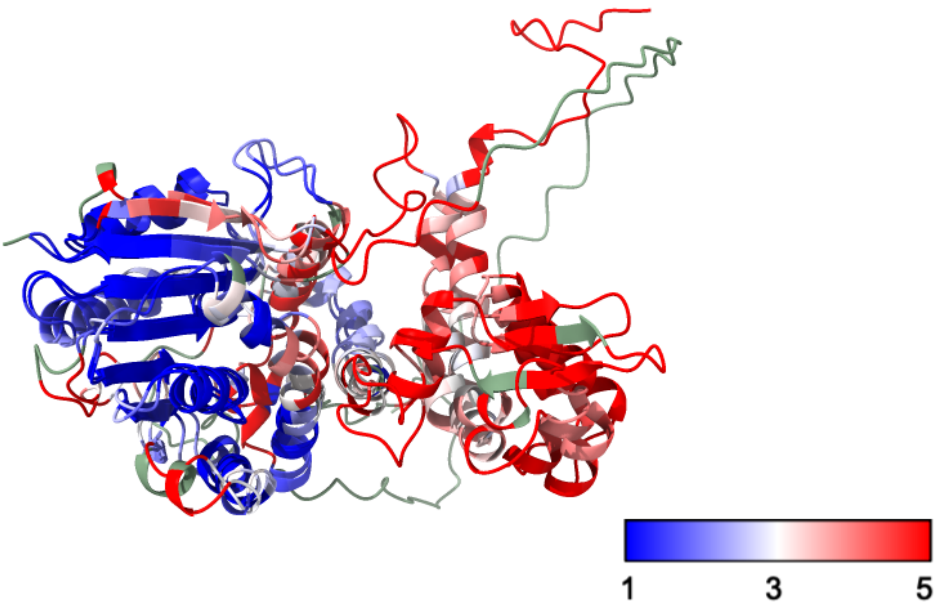


Figure SR6: ColabFold prediction of DSYB pair 2 homolog found in lucinid hosts transcriptomes in comparison to ColabFold prediction of DSYB sequence from coral *Acropora cervicornis.* Colours stated on gradient correspond to root mean square deviation in distance between aligned sequences in [Å]. No value is shown as green. The distance was calculated and visualized using ChimeraX-1.9. On the left side, the C-domain is highly conserved between these two homologs and hosts the binding sites, which supports our hypothesis of DSYB function. On the right side, in the N-domain potentially responsible for dimerization [29], the peptides show high dissimilarity.

*Partial organosulfur transformation pathways in Loripes orbiculatus symbionts suggest collaborative DMSP catabolism*

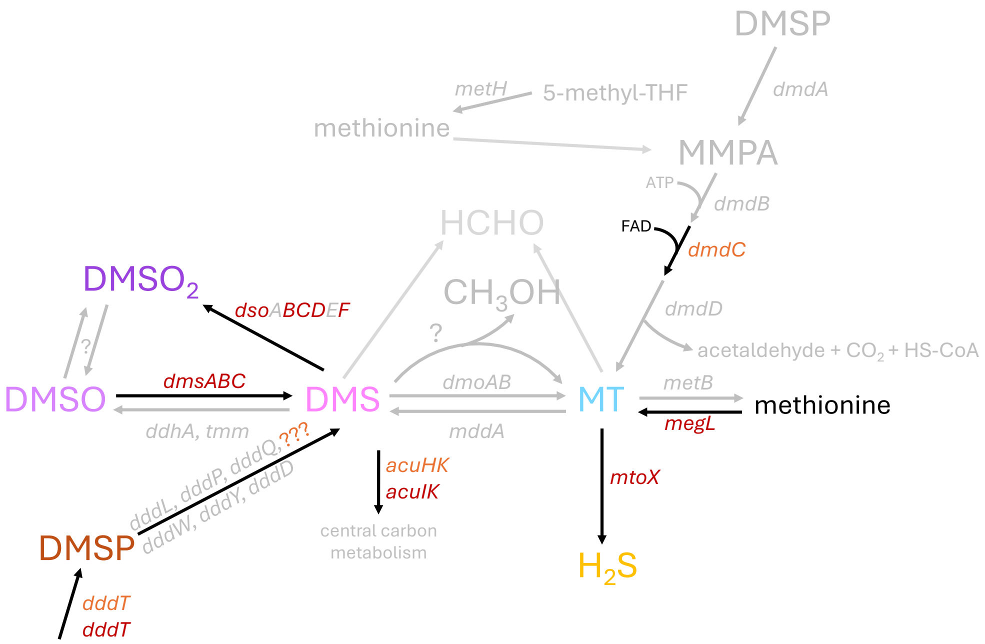


Figure SR7: Schematic representation of metabolic pathways of organosulfur compounds investigated in this study. Genes present in lucinid symbionts (*Ca.* T. luna and *Ca.* S. endoloripinus) are marked in red, and genes found in *Endozoicomonas* sp. are marked in orange. Based on Tanabe and Dahl, 2023 and Moran and Durham, 2019 [30,31].

Our search for DMSP degradation genes in the genomes of both sulphur-oxidizing symbionts were inconclusive, with only the *dddT* gene found in *Ca.* T. luna, and no hits in *Ca*. S. endoloripinus (Sudo et al. 2024; Supplementary table SR2). As there is no possibility of testing the degradation capabilities by due to lack of culturable sulphur-oxidising lucinid symbiont representatives, it is impossible to rule out the potential function. However, because there is no significant different in DMSP levels in lucinid tissues despite of differing symbiont abundances, we hypothesize that sulphur-oxidizing symbionts are not capable of DMSP catabolism, rather they use it for its anti-stress properties [32].
Another previously undescribed pathway in lucinid symbionts is re-oxidation of DMS to DMSO or $DMSO_2$. The multicomponent DMS monooxygenase DsoABCDEF catalyses the two-step oxidation of DMS to $DMSO_2$, with DMSO as an intermediate [33]. Genes encoding DsoBCDF were detected in the sulfur-oxidizing symbiont genomes of both lucinid species, whereas DsoAE were absent. While *dsoE* is considered essential for DMS oxidation, *dsoA* is not required for DMSO formation [34]. The DsoABCDEF system remains poorly characterized, and further work is needed to establish whether the identified gene complement supports DMS oxidation to $DMSO_2$ in these symbionts.

*Antibiotics and anoxic conditions increase DMSP concentrations*

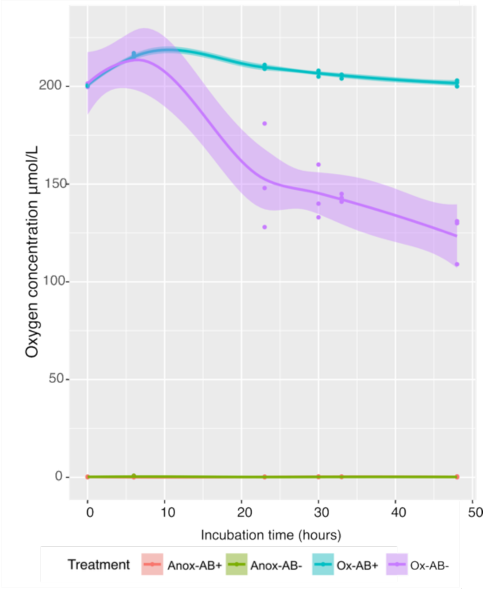


Figure SR8: Oxygen concentration in the antibiotic treatment experiment. The treatments included anoxic incubation with antibiotics (Anox-AB+) or without them (Anox-AB-) and oxic incubations with antibiotics (Ox-AB+) or without them (Ox-AB-).

DMSP is often considered a stress indicator in animals as it accumulates after exposure to direct sunlight, thermal stress, or oxidative stressors [35–37]. Despite its beneficial properties, high DMSP levels can attract pathogenic bacteria, posing risks to holobiont health [38]. To better understand the factors influencing DMSP cycling, including its production and degradation, we incubated live *Loripes orbiculatus*, the only species available for such incubation experiments, with and without antibiotics, under oxic and anoxic conditions. In the oxic controls, oxygen levels steadily declined, indicating holobiont activity (Figure SR8). However, in the oxic treatments with antibiotics, oxygen levels remained high and stable, suggesting diminished activity. All anoxic treatments remained consistently anoxic throughout the incubation. DMSP concentrations were significantly higher under anoxia compared to the oxic conditions in treatments without antibiotics ($p \leq 0.01$) (Figure 2). This suggest that anoxia-induced stress triggers increased DMSP production in *L. orbiculatus*. Moreover, antibiotic treatment resulted in a significant increase in DMSP levels in the body and gills of *L. orbiculatus* under both oxic ($p \leq 0.01$, compared to oxic controls) and anoxic ($p \leq 0.05$, compared to anoxic controls) conditions (Figure 2). These findings suggest that the lucinid microbiome plays a crucial role in regulating DMSP concentrations. Under anoxic conditions with antibiotics, DMSP concentrations were significantly higher ($p \leq 0.01$) than in oxic conditions with antibiotics, further illustrating that host stress contributes to DMSP production in the absence of microbial degradation (Figure 2). DMSP synthesis during anoxia may protect against reactive oxygen species (ROS), preventing host stress and promoting holobiont stability. While anoxia induces a stress response and DMSP production in the host, the symbionts, *Ca.* T. luna and *Ca.* S. endoloripinus, may switch to alternative electron acceptors such as DMSO. DMSO reductase was previously reported as one of the most highly expressed genes in *Loripes orbiculatus* symbionts [39], and in other sulphur-oxidising symbionts such as *Ca.* Thiosymbion oneisti [40]. However, the substrate specificity of this enzyme requires further experimental confirmation.

While DMSP production may help the host manage stress, it can also be detrimental. DMSP serves as a chemoattractant for pathogenic *Vibrio* species, which proliferate under stress and increased host susceptibility to disease [38]. In corals, members of the microbiome, such as *Endozoicomonas acroporae*, can indirectly protect their host by degrading DMSP to DMS, towards which *Vibrio* shows no chemoattraction [38,41]. We propose that in lucinid holobionts the excess DMSP could be catabolised to DMS by *Endozoicomonas*, and the products further detoxified by *Ca*. Thiodiazotropha. This function may be critical in the dynamic oxic-anoxic interface of marine sediments, the ideal habitat of lucinid holobionts, especially within seagrass meadows, where the oxygen released by seagrass plants can cause seasonal and even diurnal shifts [42]. Therefore, a healthy microbiome capable of DMSP degradation may be essential for the lucinid holobiont's health.

References:

1. Danecek, P. *et al.* Twelve years of SAMtools and BCFtools. *GigaScience* **10**, giab008 (2021).

2. Song, L. & Florea, L. Rcorrector: efficient and accurate error correction for Illumina RNA-seq reads. *GigaSci* **4**, 48 (2015).

3. Martin, M. Cutadapt removes adapter sequences from high-throughput sequencing reads. *EMBnet.journal* **17**, 10–12 (2011).

4. Kopylova, E., Noé, L. & Touzet, H. SortMeRNA: fast and accurate filtering of ribosomal RNAs in metatranscriptomic data. *Bioinformatics* **28**, 3211–3217 (2012).

5. Shen, W., Le, S., Li, Y. & Hu, F. SeqKit: A Cross-Platform and Ultrafast Toolkit for FASTA/Q File Manipulation. *PLOS ONE* **11**, e0163962 (2016).

6. Andrews, S. & others. FastQC: a quality control tool for high throughput sequence data. (2010).

7. Ewels, P., Magnusson, M., Lundin, S. & Käller, M. MultiQC: summarize analysis results for multiple tools and samples in a single report. *Bioinformatics* **32**, 3047–3048 (2016).

8. Li, H. Aligning sequence reads, clone sequences and assembly contigs with BWA-MEM. Preprint at https://doi.org/10.48550/arXiv.1303.3997 (2013).

9. Quinlan, A. R. & Hall, I. M. BEDTools: a flexible suite of utilities for comparing genomic features. *Bioinformatics* **26**, 841–842 (2010).

10. Bushnell, B. *BBMap: A Fast, Accurate, Splice-Aware Aligner*. (2014).

11. Grabherr, M. G. *et al.* Trinity: reconstructing a full-length transcriptome without a genome from RNA-Seq data. *Nat Biotechnol* **29**, 644–652 (2011).

12. Haas, B. J. *et al.* De novo transcript sequence reconstruction from RNA-Seq: reference generation and analysis with Trinity. *Nat Protoc* **8**, 10.1038/nprot.2013.084 (2013).

13. Fu, L., Niu, B., Zhu, Z., Wu, S. & Li, W. CD-HIT: accelerated for clustering the next-generation sequencing data. *Bioinformatics* **28**, 3150–3152 (2012).

14. Manni, M., Berkeley, M. R., Seppey, M., Simão, F. A. & Zdobnov, E. M. BUSCO Update: Novel and Streamlined Workflows along with Broader and Deeper Phylogenetic Coverage for Scoring of Eukaryotic, Prokaryotic, and Viral Genomes. *Molecular Biology and Evolution* **38**, 4647–4654 (2021).

15. Blum, M. *et al.* InterPro: the protein sequence classification resource in 2025. *Nucleic Acids Research* **53**, D444–D456 (2025).

16. Mirdita, M. *et al.* ColabFold: making protein folding accessible to all. *Nat Methods* **19**, 679–682 (2022).

17. Meng, E. C. *et al.* UCSF ChimeraX : Tools for structure building and analysis. *Protein Science* **32**, e4792 (2023).

18. Pruesse, E. *et al.* SILVA: a comprehensive online resource for quality checked and aligned ribosomal RNA sequence data compatible with ARB. *Nucleic Acids Research* **35**, 7188–7196 (2007).

19. Pruesse, E., Peplies, J. & Glöckner, F. O. SINA: Accurate high-throughput multiple sequence alignment of ribosomal RNA genes. *Bioinformatics* **28**, 1823–1829 (2012).

20. Chiou, Y.-J. *et al.* Similar but different: Characterization of dddD gene–mediated DMSP metabolism among coral-associated Endozoicomonas. *Science Advances* **9**, eadk1910 (2023).

21. Lim, S. J. *et al.* Extensive Thioautotrophic Gill Endosymbiont Diversity within a Single *Ctena orbiculata* (Bivalvia: Lucinidae) Population and Implications for Defining Host-Symbiont Specificity and Species Recognition. *mSystems* **4**, 1–19 (2019).

22. Hill, R. W. & Dacey, J. W. H. Processing of ingested dimethylsulfoniopropionate by mussels Mytilus edulis and scallops Argopecten irradians. *Marine Ecology Progress Series* **343**, 131–140 (2007).

23. Hill, R. W. & Dacey, J. W. H. Exceptional accumulation and retention of dimethylsulfoniopropionate by molluscs. *Environ. Chem.* **13**, 231–238 (2015).

24. Kurahashi, M. & Yokota, A. Endozoicomonas elysicola gen. nov., sp. nov., a γ-proteobacterium isolated from the sea slug Elysia ornata. *Systematic and Applied Microbiology* **30**, 202–206 (2007).

25. Mauss, M. Endosymbionts of Two Species of Mediterranean Lucinid Clams: A Molecular, Microbial and Histological Analysis. *Uni Wien* (2008).

26. Athamena, A. *et al.* Salinity regulates N-methylation of phosphatidylethanolamine in euryhaline crustaceans hepatopancreas and exchange of newly-formed phosphatidylcholine with hemolymph. *J Comp Physiol B* **181**, 731–740 (2011).

27. Athamena, A., Trajkovic-Bodennec, S., Brichon, G., Zwingelstein, G. & Bodennec, J. Synthesis of Phosphatidylcholine Through Phosphatidylethanolamine N-Methylation in Tissues of the Mussel Mytilus galloprovincialis. *Lipids* **46**, 1141–1154 (2011).

28. Péqueux, A. Osmotic Regulation in Crustaceans. *Journal of Crustacean Biology* **15**, 1–60 (1995).

29. Li, C. *et al.* Mechanistic insights into the key marine dimethylsulfoniopropionate synthesis enzyme DsyB/DSYB. *mLife* **1**, 114–130 (2022).

30. Tanabe, T. S. & Dahl, C. HMSS2 : An advanced tool for the analysis of sulphur metabolism, including organosulphur compound transformation, in genome and metagenome assemblies. *Molecular Ecology Resources* 1755–0998.13848 (2023).

31. Moran, M. A. & Durham, B. P. Sulfur metabolites in the pelagic ocean. *Nature Reviews Microbiology* **17**, 665–678 (2019).
32. Kiene, R. P., Linn, L. J. & Bruton, J. A. New and important roles for DMSP in marine microbial communities. *Journal of Sea Research* **43**, 209–224 (2000).
33. Boden, R. & Hutt, L. P. Bacterial Metabolism of C1 Sulfur Compounds. in *Aerobic Utilization of Hydrocarbons, Oils and Lipids* (ed. Rojo, F.) 1–43 (Springer International Publishing, Cham, 2019).
34. Horinouchi, M., Yoshida, T., Nojiri, H., Yamane, H. & Omori, T. Polypeptide Requirement of Multicomponent Monooxygenase DsoABCDEF for Dimethyl Sulfide Oxidizing Activity. *Bioscience, Biotechnology, and Biochemistry* **63**, 1765–1771 (1999).
35. Raina, J.-B. *et al.* DMSP biosynthesis by an animal and its role in coral thermal stress response. *Nature* **502**, 677–680 (2013).
36. Gardner, S. G. *et al.* Dimethylsulfoniopropionate, superoxide dismutase and glutathione as stress response indicators in three corals under short-term hyposalinity stress. *Proceedings of the Royal Society B: Biological Sciences* **283**, 20152418 (2016).
37. Deschaseaux, E. S. M. *et al.* Effects of environmental factors on dimethylated sulfur compounds and their potential role in the antioxidant system of the coral holobiont. *Limnology and Oceanography* **59**, 758–768 (2014).
38. Garren, M. *et al.* A bacterial pathogen uses dimethylsulfoniopropionate as a cue to target heat-stressed corals. *ISME J* **8**, 999–1007 (2014).
39. Petersen, J. M. *et al.* Chemosynthetic symbionts of marine invertebrate animals are capable of nitrogen fixation. *Nat Microbiol* **2**, 16195 (2016).
40. Paredes, G. F. *et al.* Anaerobic Sulfur Oxidation Underlies Adaptation of a Chemosynthetic Symbiont to Oxic-Anoxic Interfaces. *mSystems* **6**, 10.1128/msystems.01186-20 (2021).
41. Tandon, K. *et al.* Comparative genomics: Dominant coral-bacterium Endozoicomonas acroporae metabolizes dimethylsulfoniopropionate (DMSP). *ISME J* **14**, 1290–1303 (2020).
42. Lee, K.-S. & Dunton, K. H. Diurnal changes in pore water sulfide concentrations in the seagrass Thalassia testudinum beds: the effects of seagrasses on sulfide dynamics. *Journal of Experimental Marine Biology and Ecology* **255**, 201–214 (2000).
